# A unique chaperoning mechanism in Class A JDPs recognizes and stabilizes mutant p53

**DOI:** 10.1101/2023.11.28.568993

**Authors:** Guy Zoltsman, Thi Lieu Dang, Miriam Kuchersky, Ofrah Faust, Micael S. Silva, Tal Ilani, Anne S. Wentink, Bernd Bukau, Rina Rosenzweig

**Affiliations:** Department of Chemical and Structural Biology, Weizmann Institute of Science, Rehovot, 761000, Israel; Center for Molecular Biology of Heidelberg University (ZMBH) and German Cancer Research Center (DKFZ), DKFZ-ZMBH Alliance, Im Neuenheimer Feld 282, Heidelberg D-69120, Germany; Leiden Institute of Chemistry, Leiden University, Einsteinweg 55, 2333CC Leiden, The Netherlands

**Keywords:** Molecular chaperones, protein misfolding, NMR, J-domain proteins, protein aggregation, p53

## Abstract

J-domain proteins (JDPs) constitute a large family of molecular chaperones that bind a broad spectrum of substrates, targeting them to Hsp70, thus determining the specificity and activating the entire chaperone functional cycle. The malfunction of JDPs is therefore inextricably linked to myriad human disorders. Here we uncover a novel mechanism by which chaperones recognize misfolded clients, present in class-A JDPs. Through a newly-identified β-hairpin site, these chaperones detect changes in protein dynamics at the initial stages of misfolding, prior to exposure of hydrophobic regions or large structural rearrangements. The JDPs then sequester misfolding-prone proteins into large oligomeric assemblies, protecting them from aggregation. Through this mechanism, class-A JDPs bind destabilized p53 mutants, preventing clearance of these oncoproteins by Hsp70-mediated degradation, thus promoting cancer progression. Removal of the β-hairpin abrogates this protective activity while minimally affecting other chaperoning functions. This suggests the class-A JDP β-hairpin as a highly specific target for cancer therapeutics.

## INTRODUCTION

Molecular chaperones are a diverse group of proteins crucial for maintaining protein homeostasis and protecting cells against stress ^1^. Improper function of these essential machines can lead to the accumulation of misfolded and aggregated proteins, which have been correlated with a wide array of human disorders, including neurodegeneration, myopathies, and cancer ^2–5^.

The Hsp70 system acts in the regulation of protein function and as the first line of defense against misfolding/proteotoxic stress by facilitating the refolding and aggregation-prevention of misfolded proteins, and the transfer of damaged proteins for degradation ^1,6^. These crucial activities of Hsp70s are tightly regulated by J-domain proteins (JDPs), that not only pre-select and deliver non-natively folded proteins to the chaperones, but also activate these machines by stimulating Hsp70 ATP hydrolysis, inducing stable substrate binding ^7^. Recently, several studies have indicated that JDPs can also function independently as bona fide chaperones, utilizing holdase activity to prevent the aggregation of their client proteins ^8, 9^. In humans, JDPs constitute the largest co-chaperone family, with nearly 50 different paralogs, which vary in their structures and substrate selectivity ^4,10^. However, despite playing key roles in a myriad of cellular activities, how these chaperones recognize and affect non-natively folded clients in the cell remains poorly understood.

The p53 tumor suppressor protein is a known cellular client of the human Hsp70 chaperone system, which regulates its conformational stability and prevents p53 ubiquitination and degradation ^11,12^. Increasing evidence indicates direct involvement of different JDP family members in cancer, with both tumor-suppressive and oncogenic roles reported ^3,13,14^. Specifically, several cancer-associated conformational (destabilized) p53 mutants have been shown to interact with JDP chaperones ^3^.

In this work, we use such destabilized oncogenic p53 mutants as representative model substrates to obtain a structural and mechanistic understanding of the interaction of JDP chaperones with conformationally unstable clients. We find that only a specific subset of JDPs, class A, can recognize and bind to the p53 mutants. Using nuclear magnetic resonance (NMR) spectroscopy, we further show that the chaperones utilize a previously uncharacterized β-hairpin client-binding site to detect the initial misfolding in the DNA-binding domain of p53 and stabilize the mutant protein. Excitingly, we find that class A JDPs recognize such destabilization via a unique mechanism not reported for any other chaperones - sensing the increased dynamics and transient breakage of hydrogen bonds associated with the early stages of misfolding. Moreover, once bound, class A JDPs form large oligomeric structures with these clients, sequestering and shielding them from aggregation during stress. Such a sequestration mechanism has not been described for any other chaperone in the Hsp70 system, and allows class A JDPs to pre-emptively protect β-sheet rich proteins on the verge of becoming misfolded. This, therefore, represents a novel functional mode by which the protein quality control system can efficiently counteract the deleterious misfolding of β-sheet proteins. However, in the context of cancer, this chaperone stabilization of p53 oncoproteins prevents their removal by cellular quality control machineries, effectively promoting cancer cell survival and disease progression.

## RESULTS

### A specific JDP class prevents aggregation of p53 hot-spot mutants

Of specific interest are oncogenic p53 mutations that cause destabilization of the p53 DNA-binding domain (DBD) structure and account for ∼40% of cancer-associated p53 mutants ^15–19^. These mutant p53s usually have wild-type-like structures at lower temperatures, but undergo misfolding and rapid aggregation at body temperature ^12, 20, 21^. As such non-native, aggregation-prone folds are typical clients of molecular chaperones, we screened for JDPs that specifically recognize and then prevent the misfolding and aggregation of two destabilizing, cancer-associated hot-spot p53 mutants - R249S and R282W. Four main classes of JDPs, previously reported to affect cancer progression, were tested: class A (DNAJA1 and DNAJA2), class B (DNAJB1 and DNAJB4), non-canonical class B (DNAJB2 and DNAJB6), and class C (DNAJC7 and DNAJC8) chaperones ^3, 13, 14^.

At physiological temperatures, the R249S and R282W p53 mutants misfold and aggregate rapidly, as evident from the increase in light scattering (Figures 1A-C). In the presence of class A JDPs, this aggregation was significantly inhibited, with DNAJA1 increasing the half-time of p53 aggregation from 30 to 55 minutes, and DNAJA2 chaperone completely suppressing mutant p53 aggregation for over 8 hours (Figures 1A,C). DNAJA2 was further found to significantly increase the amounts of mutant p53 found in the soluble fraction, with only 2% and 4% of R249S and R282W, respectively, being detected in the insoluble pellet fraction after 5 hours at 37°C (Figure S1A). In the absence of DNAJA2 nearly all p53 mutant protein was found in the pellet (Figure S1A). No such aggregation-inhibiting activity was detected, however, for any of the class B or class C JDPs tested (Figures 1A-C).

**Figure 1.**
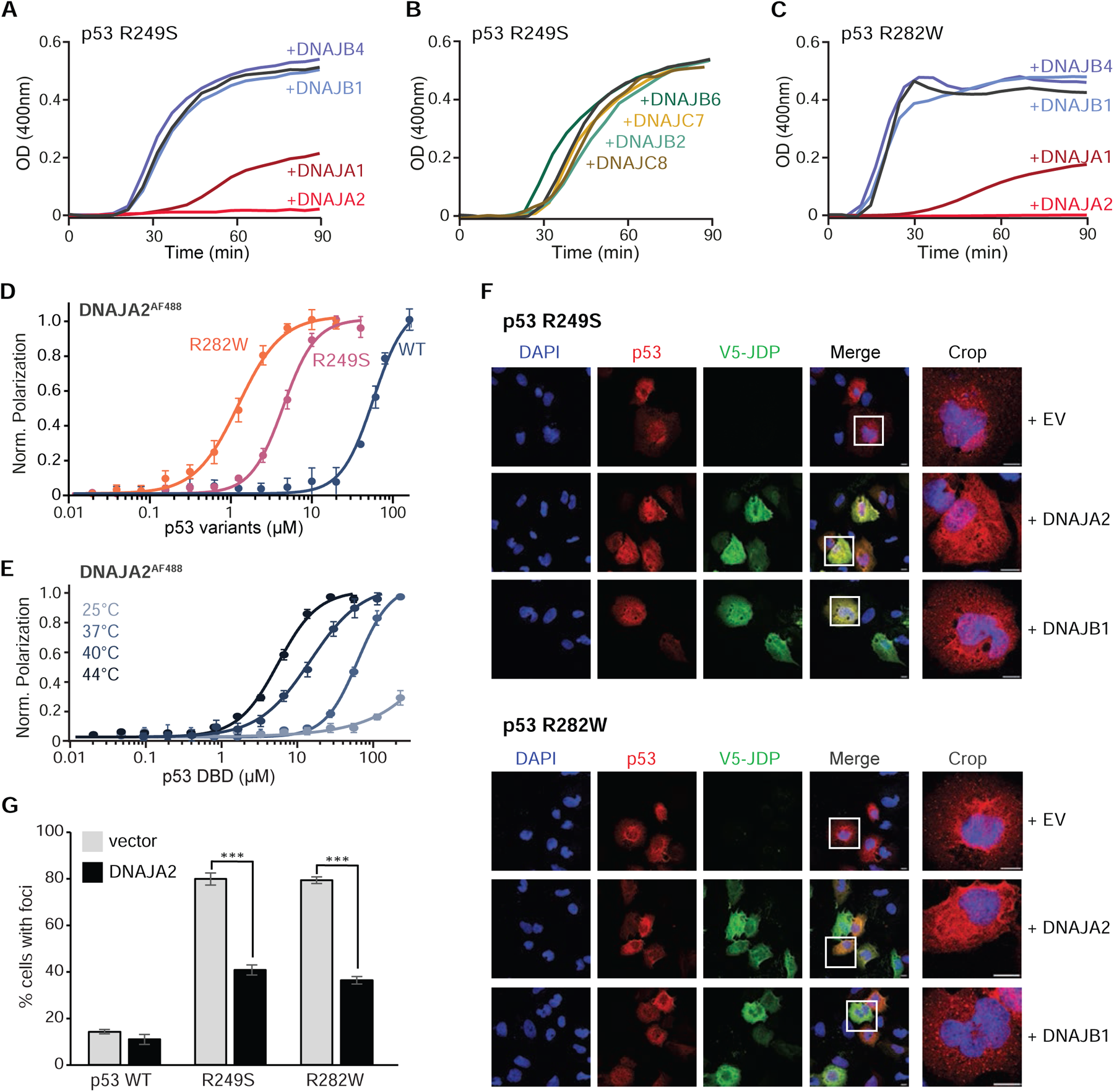
Class A JDPs prevent misfolding and aggregation of mutant p53. **(A-B)** Aggregation of R249S destabilized p53 mutant alone (black), or upon addition of 2-fold molar excess of DNAJA1 (maroon), DNAJA2 (red), DNAJB1 (blue), DNAJB4 (purple), DNAJB2 (teal), DNAJB6 (green), DNAJC7 (yellow), or DNAJC8 (brown), monitored by light scattering. Data are means (n = 3). **(C)** Aggregation of R282W destabilized p53 mutant alone (black), or in the presence of DNAJA1 (maroon), DNAJA2 (red), DNAJB1 (blue), or DNAJB4 (purple) JDPs monitored by light scattering. Data are means (n = 3). **(D)** Fluorescence anisotropy binding assays of Alexa Fluor-488 labeled DNAJA2, titrated with increasing concentrations of p53 WT (blue), R249S (pink), or R282W (orange), measured at 37°C. R249S and R282W p53 mutants bind DNAJA2 with 4.4±0.2 μM and 1.2±0.1 μM affinity, while only a weak binding is detected for WT p53. Data are means ± SEM (n = 3). **(E)** Fluorescence anisotropy binding assays of Alexa Fluor-488 labeled DNAJA2 titrated with increasing concentrations of p53 WT at increasing temperatures (25-44°C; light to dark blue). DNAJA2 affinity for WT p53 increased with temperature; from ∼70 μM at 37°C to 10.4 μM at 40°C, and 4.5 μM at 44°C. Data are means ± SEM (n=3). **(F)** Immunofluorescence staining of destabilized R249S (top) or R282W (bottom) p53 mutant overexpressed in the p53 null SaOS2 cell lines shows distinct accumulation of p53 in cytoplasmic foci, corresponding to aggregates ^20^. Co-expression of V5-DNAJA2, but not V5-DNAJB1, results in a significant increase in diffuse cytoplasmic staining of mutant p53. EV - empty vector. The crop images are the overlay of p53 signal with DAPI. Scale bar: 10 μm. **(G)** Quantification of cytoplasmic foci in cells overexpressing p53 WT, R249S or R282W with and without DNAJA2 co-expression (representative images in Figures 1F and S1F). Data represents mean values ± s.d (n=3). *** p<0.001 (Student’s *t*-test). See also Figure S1.

NMR binding experiments further established that only class A JDPs interact with the mutant p53, while other JDPs, such as DNAJB1 do not (Figures S1B-D). Thus, the class-specific JDP chaperoning activity is directly linked to the ability of the chaperones to recognize and bind the mutant p53.

p53 R249S and R282W mutants are thermodynamically unstable proteins ^22, 23^, with DBD melting temperatures of 37.3±0.1°C and 35.3±0.6°C, respectively (Figure S1E). Thus, at physiological temperature R249S and R282W are largely destabilized, while the WT p53 protein, which has a melting temperature of 42.9±0.1°C, remains folded (>95%) ^24^ (Figure S1E). To test if class A JDPs can specifically recognize the destabilized / misfolded conformations of p53, we used fluorescence anisotropy to determine the dissociation constants of Alexa Fluor 488-labeled DNAJA2 to both WT p53 and p53 mutants. DNAJA2 bound to R249S and R282W with 4.4 ± 0.2 μM and 1.2 ± 0.1 μM affinities, respectively. In contrast, the interaction with the WT p53 protein was very weak, and saturation of binding was not achieved over the sampled concentration range (Figure 1D).

In order to validate that the chaperones indeed recognize p53 destabilization and not a unique fold or sequence motif in the p53 hot-spot mutants, we measured the affinities of DNAJA2 towards WT p53 at elevated temperatures (40-44°C), which trigger WT p53 misfolding ^23^ (Figure 1E). The affinity of DNAJA2 to WT p53 increased significantly at elevated temperatures, and at 44°C was comparable to those measured for the hot-spot p53 mutants (4.5 ± 0.2 μM, Figure 1E).

Thus, class A chaperones are unique amongst JDPs in that they can prevent the aggregation of mutant p53 species by specifically recognizing the non-native, destabilized conformations of these proteins.

### Class A JDPs prevent p53 aggregation in cells

To investigate whether the observed chaperoning activity of class A JDPs translates to the cellular environment, we tested the effect of JDP overexpression in a cell culture model of mutant p53 aggregation ^20^. Transient transfection of SaOS2 cells (which lack endogenous p53) with plasmids encoding the R249S and R282W mutants resulted in an accumulation of p53 in cytoplasmic “foci” in 80% of cells (Figures 1F, G). In contrast, wild type p53 was found predominantly in the nucleus, and cytoplasmic foci were observed in only 14% of cells (Figure 1G and S1F). Blue-Native PAGE, followed by western blot analysis, confirmed that while WT p53 existed predominantly in a native tetramer state, p53 R249S and R282W were present in large aggregates (> 200 kDa) (Figure S1G), in agreement with previous reports ^20^.

The simultaneous overexpression of mutant p53 with class A DNAJA2 significantly reduced the percentage of cells with visible cytoplasmic foci (to 45% for R249S and 39% for R282W), with no such effect observed for the class B JDP DNAJB1 (Figures 1F-G and S1F,H). Likewise, in differential centrifugation experiments, the amount of p53 found in the insoluble pellet fraction was reduced by 52% and 45% for R249S and R282W respectively in the presence of DNAJA2 (Figures S1I-J).

Combined, these observations establish a class-specific aggregation-prevention activity for class A JDPs on destabilized p53 mutants in human cells.

### Characterizing the interaction of DNAJA2 chaperone with p53 DBD

To elucidate how class A JDPs specifically recognize the destabilized p53 conformers, we obtained structural information for the DNAJA2-p53 complex using NMR (Figures 2A and S2A-B). Given the rapid precipitation of mutant p53s at 37°C, binding experiments were conducted at 28°C for R249S and R282W variants of p53 DBD. Under these conditions, the mutants aggregated significantly slower, while still exhibiting prominent binding to DNAJA2 with affinities of 6.3±0.7 (R282W) and 55.9±2.6 (R249S) µM (Figure S2C).

**Figure 2.**
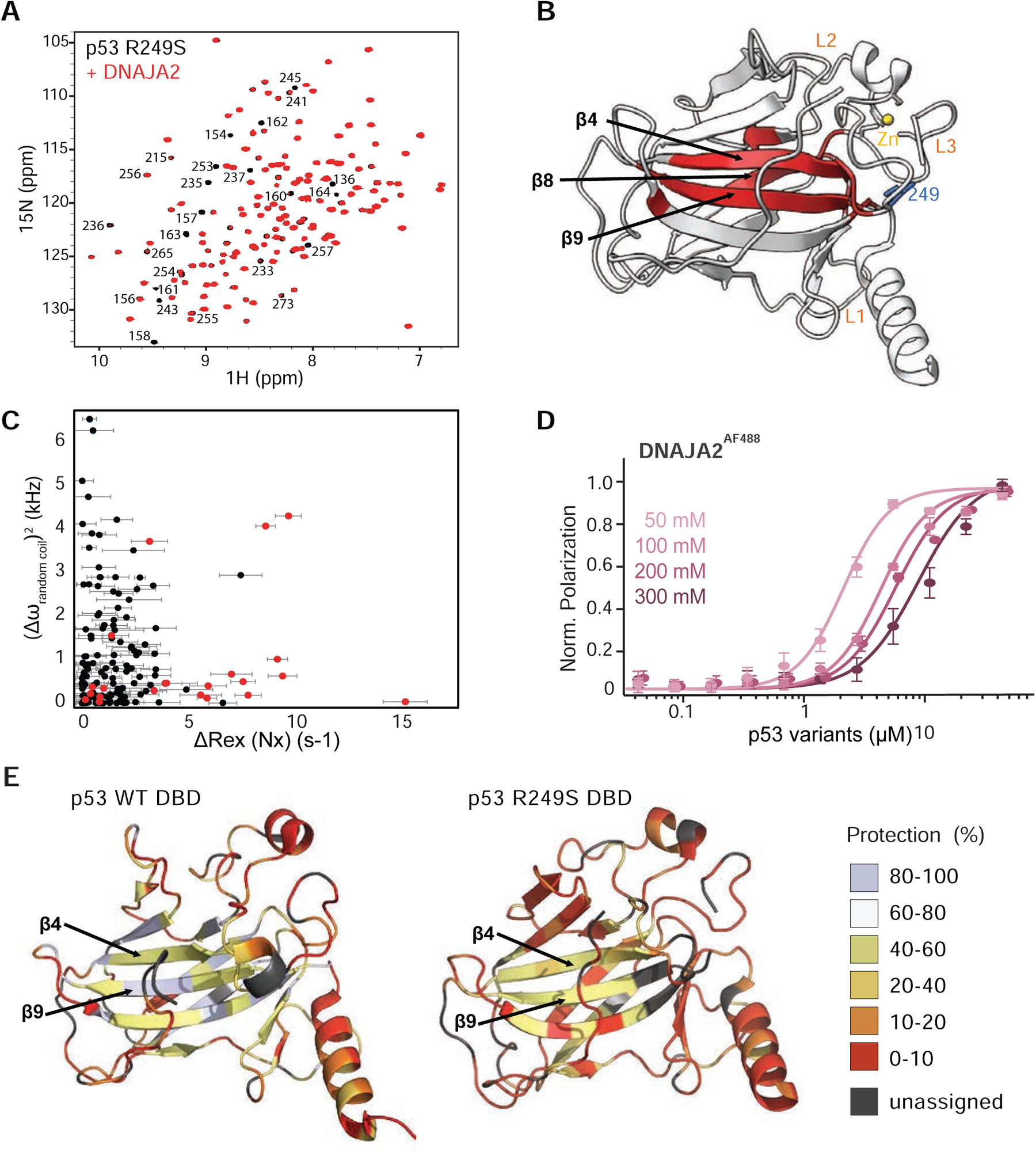
Mapping the interaction of p53 R249S mutant with DNAJA2 chaperone. **(A)** Overlay of two-dimensional ^1^H - ^15^N HSQC spectra of ^15^N, ^2^H labeled R249S DBD alone (black), and with 2-fold molar excess of deuterated (^2^H) DNAJA2 (red). **(B)** Cartoon representation of p53 R249S structure (PDB: 3D07), with residues showing significant binding to DNAJA2 (I/I_0_<0.4) colored red. **(C)** Microsecond exchange contributions (R_ex,µs_) to ^15^N transverse relaxation rates of DNAJA2-bound p53 obtained from measurement of residue-specific R_2_(2H_x_N_z_), R_2_(2H_z_N_x_), R_2_(2H_x_N_x_), and R_1_(2H_z_N_z_) relaxation rates (which should correlate with changes in chemical shifts between the bound and free p53 states ^37, 38^) plotted as a function of changes in chemical shifts between free p53 and unfolded, random coil ^79–81^, values (Δϖ_random coil_). p53 residues affected by DNAJA2 binding are colored red. Lack of correlation indicates that the DNAJA2-bound p53 is not found in an unfolded conformation. **(D)** Fluorescence anisotropy binding assays of Alexa Fluor-488 labeled DNAJA2 titrated with increasing concentrations of p53 R249S at increasing NaCl concentrations (50-300 mM; light to dark pink). DNAJA2 affinity for p53 R249S decreased with increase in ionic strength indicative of electrostatic interaction. Data are means ± SEM (n=3). **(E)** Degree of solvent protection / accessibility, calculated based on H/D exchange experiments, for WT (left) and R249S (right) p53 DBD at 28°C, colored on their corresponding crystal structures (PDB; WT:2OCJ, R249S: 3D07). See also Figure S2 and S3.

Upon addition of deuterated (^2^H) DNAJA2 to ^2^H,^15^N-labeled p53 R249S or R282W DBDs, significant peak broadening occurred in p53 residues 155-163 (β4), 232-237 (β8) and 250-259 (β9) (Figures 2A-B and S2A-B), indicative of interaction with DNAJA2 occurring at an intermediate NMR time-scale. Notably, the regions in the mutant p53s which were recognized by DNAJA2 were structured β-strands that form the protein core. However, the ^1^H-^15^N heteronuclear single quantum coherence (HSQC) spectrum of p53 R249S DBD showed no major differences between the free and the DNAJA2-bound states, indicating that upon interaction with the chaperone, the global fold of the β-rich core region remains intact (Figure 2A). Thus, unlike Hsp70 chaperones ^25, 26^, class A JDP binding does not cause global or partial unfolding of p53, but rather maintains p53 mutants in a folded or nearly-folded conformation.

Intrigued by this unusual mode of interaction with structured elements, we aimed to gain a detailed structural characterization of the chaperone-bound conformation of the p53 mutant. However, since the binding between mutant p53 and DNAJA2 occurs on a µs-ms time-scale, causing peak-broadening, structural information for DNAJA2-bound p53 cannot be obtained via direct NMR measurements. In such cases, it is possible to derive this information indirectly, utilizing chemical exchange between the free and bound states, by detecting the visible NMR peaks of the free p53 ^27–30^. Such an approach requires a sample where only a small population of p53 is in complex with the chaperone. Accordingly, we measured a series of NMR dynamic experiments covering different µs-ms timescale ranges on a sample containing 15% p53-DNAJA2 complex. No minor conformations indicative of DNAJA2-binding were detected in ^15^N Chemical Exchange Saturation Transfer (CEST) ^31, 32^ and ^15^N Carr-Purcell-Meiboom-Gill (CPMG) relaxation dispersion ^33, 34^ NMR experiments (Figure S2D). As these experiments are sensitive to kinetic processes on the millisecond time scale ^35^, we concluded that p53 binding to DNAJA2 occurs at a faster rate of microseconds (µs).

We then recorded NMR experiments measuring the decay rates of four coherences [R_2_(2H_x_N_z_), R_2_(2H_z_N_x_), R_2_(2H_x_N_x_), R_1_(2H_z_N_z_)] ^36, 37^, which provide site-specific values for exchange contributions on the µs time scale (R_ex,µs_). Such isolation is possible since contributions from processes in the ms timescale (*k*_ex_ < 2000 s^-1^) are effectively suppressed when relaxation rates are measured under spin-locking conditions ^36, 37^. These experiments showed a clear DNAJA2-dependent increase in R_ex,µs_ rates in residues 155-163 (β4) and 250-259 (β9) of p53 R249S, which correspond to DNAJA2 binding sites on p53 (Figures S2E-F). We next plotted this increase in R_ex,µs_ (which is proportional to the chemical shift differences between the DNAJA2-bound and free p53 states ^38^) as a function of chemical shift differences between free and random-coil shifts of unfolded p53 (see materials and methods for more information). No such correlation was detected (Figure 2C), confirming that DNAJA2 does not recognize the unfolded, but rather the nearly folded state in the destabilized p53 mutants.

Such a mode of interaction with structured, nearly native folded elements is quite unusual for molecular chaperones, which typically recognize exposed and unstructured hydrophobic regions in their client proteins ^39,40^. As the DBD core of p53, and in particular the β9 region are enriched in hydrophobic amino acids, we next asked if DNAJA2 is similar to other canonical chaperones in that it, too, recognizes its clients through hydrophobic interactions. We thus determined the affinity of the chaperone for p53 R249S upon increased ionic strength, which should increase hydrophobic attraction. Surprisingly, we found that the affinity of p53 for DNAJA2 in fact decreased 4-fold between 50 mM (1.9±0.1 μM) and 300 mM NaCl (8.0±0.6 μM) (Figure 2D), indicating that this binding is of an electrostatic nature and not governed by hydrophobic interactions.

DNAJA2 thus recognizes the destabilized p53 via a new and unusual chaperoning mechanism - by binding to structured regions in misfolding-prone, destabilized, but nearly-folded clients via electrostatic interactions. This is in stark contrast to canonical chaperones, that typically bind exposed, unstructured hydrophobic regions in their client proteins ^39, 40^.

### DNAJA2 chaperones sense the transient breakage of hydrogen bonds in p53

As DNAJA2 chaperones recognize nearly-folded clients, this raises the question of how these class A JDPs can discriminate between properly folded and destabilized p53 mutants. While, structurally, p53 WT and R249S and R282W mutant DBDs are highly similar, several studies have attributed the decrease in stability of the mutants to an increase in the mobility of their core β-sheets, eventually leading to unfolding and protein aggregation ^41–44^.

To test whether an increase in local structural mobility is what exposes the DNAJA2 chaperone-binding sites in p53 hot-spot mutants, we recorded a series of hydrogen-deuterium exchange NMR measurements. H/D-exchange reports on the ability of protein backbone amides to exchange hydrogen with deuterium presented as D_2_O. Residues that are in unstructured or solvent-exposed regions exchange rapidly (seconds-minutes), while amide protons in structured and hydrogen-bonded positions exchange slowly (hours-days), and are described as “protected”.

H/D-exchange time courses were collected for WT p53 and R249S mutant DBDs. The overall exchange profiles for the WT protein correlated well with the presence of secondary structure elements, with β1-10 showing high degrees of protection, while the rest of the protein was highly dynamic, in agreement with previous reports ^26^ (Figure 2E). Notable differences, however, were observed between the WT and the destabilized p53 R249S mutant, with substantially decreased protection in β1, β2, β5-7, β8, and β10 (Figures 2E and S3A-B). β4 and β9 residues, that form the main chaperone-binding interface, showed only a moderate, ∼25% decrease in protection in p53 R249S (Figures 2E and S3A-B). Interestingly, these regions, which are the last to be exposed in the destabilized mutants, were previously identified as the main aggregation-prone sequence of p53 ^20, 45^ and are the primary site of interaction with DNAJA2.

Thus, hot-spot p53 oncogenic mutations cause destabilization of the hydrogen bond network and increased dynamics in the DBD, exposing the aggregation-prone β4 and β9 strands. Similar destabilization and core exposure was also measured for the WT p53 at elevated temperatures (Figure S3C). Such exposure can lead to misfolding and subsequent aggregation via the β9 strand, but likewise increases the availability for recognition and binding by DNAJA2. This chaperone binding can then protect these dynamic regions via direct physical interaction, stabilizing the DBD fold, thereby preventing further misfolding and aggregation.

Class A JDPs thus emerge as a new type of molecular chaperone, capable of recognizing and preventing the initial stages of protein misfolding in their clients by sensing the transient breakage of hydrogen bonds.

### DNAJA2 binds p53 through a new client binding site

There is currently only a limited structural and mechanistic understanding of how JDPs recognize and bind their clients. In the case of class A JDPs, C-terminal domains I and II (CTDI and CTDII) were shown to be the binding sites for unfolded polypeptides ^46–48^, with the zinc-finger-like region (ZFLR) and the disordered G/F-rich domain (GF) potentially also contributing to this interaction ^47, 49, 50^ (Figure 3A). To date, however, no structural studies have been published showing JDP interactions with either partially- or non-natively folded (misfolded) proteins.

**Figure 3.**
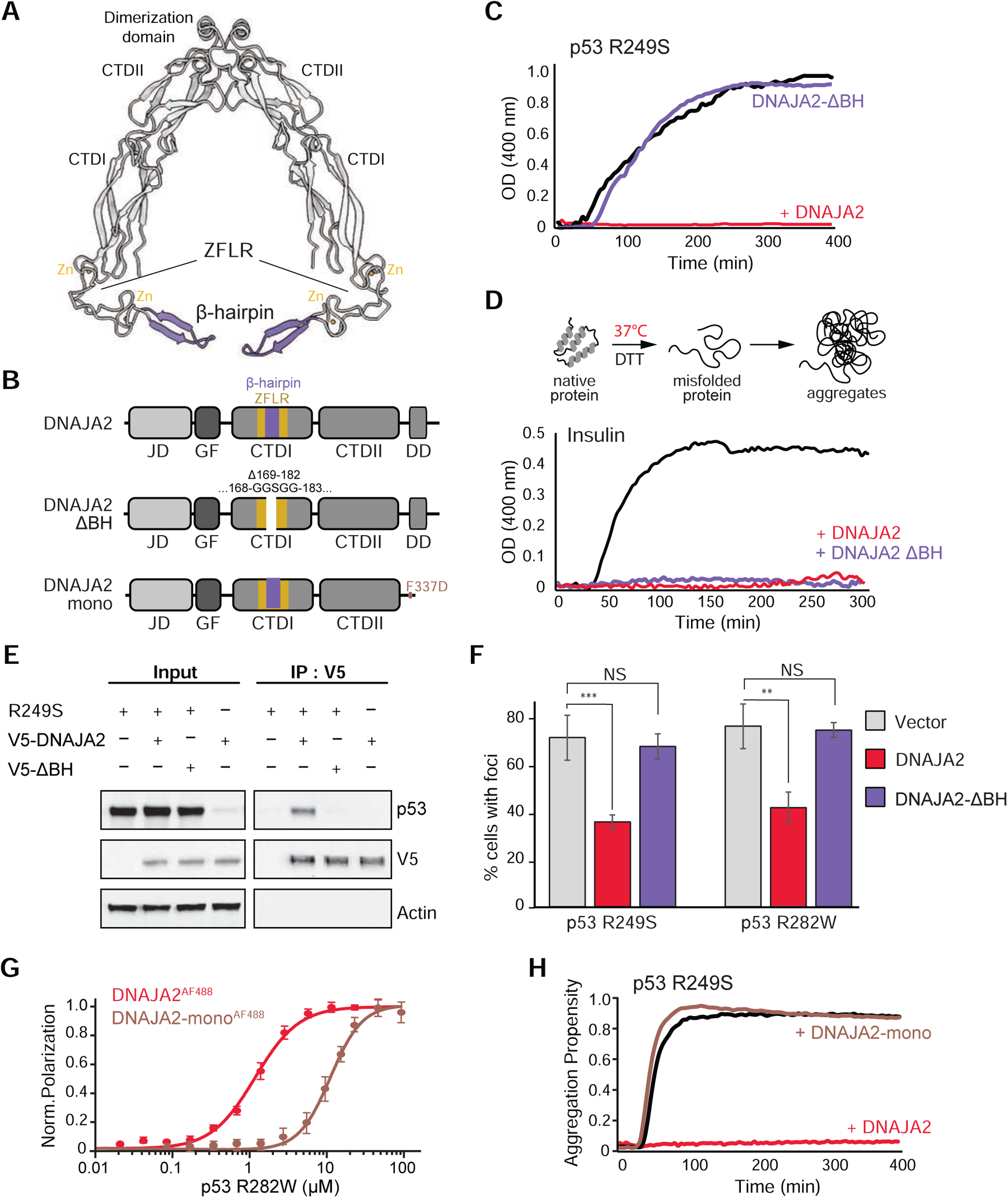
DNAJA2 recognizes misfolded p53 by the β-hairpin insertion in the ZFLR. **(A)** Cartoon representation of DNAJA2 structural model, with the residues identified by NMR experiments to interact with p53 (Figures S3A-B) colored in violet. p53 binds to a β-hairpin insertion in the zinc-finger-like region (ZFLR). **(B)** Domain organization of wild-type DNAJA2 (top), DNAJA2 mutant lacking the β-hairpin (DNAJA2^ΔBH^; bottom), and a truncated monomeric DNAJA2 construct lacking the dimerization domain (DNAJA2^mono^; bottom). The zinc finger-like region (ZFLR) is colored yellow, the β-hairpin in violet, and the Δ169-182 deletion and its substitution with GGSGG sequence is represented as a gap in the ZFLR. **(C)** Light scattering measurements (OD 400 nm) following aggregation of R249S alone (black), or in the presence of WT DNAJA2 (red) or DNAJA2^ΔBH^ mutant (violet). No aggregation-prevention activity was detected for the mutant DNAJA2 lacking the β-hairpin region. Data are means (n = 3). **(D)** Light scattering measurements (OD 400 nm) following aggregation of heat-denatured insulin alone (black), or in the presence of 3-fold molar excess of WT DNAJA2 (red) or DNAJA2^ΔBH^ mutant (violet). Both DNAJA2 WT and DNAJA2^ΔBH^ mutant fully suppress insulin aggregation over 5 hours. Data are means (n = 3). **(E)** Co-immunoprecipitation experiment of R249S p53 with either WT DNAJA2 or DNAJA2^ΔBH^ mutant tagged with V5 from SaOS2 cell lysates. Mutant p53 co-precipitated only with WT DNAJA2 but not the DNAJA2^ΔBH^ mutant. **(F)** Quantification of cells with cytoplasmic foci in SaOS2 cell lines overexpressing p53 mutants (R249S, R282W) and upon co-expression of V5-DNAJA2 WT (red) or V5-DNAJA2^ΔBH^ (violet). The DNAJA2 β-hairpin region is required to prevent mutant p53 aggregation in cells. Data represent mean values ± s.d (n=3). ** p<0.01, *** p<0.001, NS not significant (one-way ANOVA with Bonferroni’s multiple comparison test). **(G)** Fluorescence anisotropy binding assays of Alexa Fluor-488 labeled DNAJA2 WT dimer (red) and DNAJA2 monomer (brown), titrated with increasing concentrations of R282W p53 mutant. Disruption of the DNAJA2 dimerization domain weakens the binding, increasing the K_D_ from 1.0±0.1 to 5.1±0.2 µM for p53 R282W DBD. Data are means ± SEM (n = 3). **(H)** Aggregation of R249S destabilized p53 mutant alone (black), and upon addition of 2-fold molar excess of either DNAJA2 WT dimer (red) or DNAJA2 monomer (brown). Only the dimeric DNAJA2 can prevent p53 aggregation, while the monomeric version of the chaperone shows no activity. Data are means (n = 3). See also Figures S4 and S5.

To elucidate which structural regions in DNAJA2 interact with the destabilized forms of p53, we recorded the ^1^H-^15^N HSQC spectra of a ^2^H, ^15^N-labeled monomeric version of DNAJA2 upon addition of the two destabilized p53 mutants. As no chemical shift assignments are currently available for the human DNAJA2, the NMR measurements were performed on the yeast homologue, Ydj1.

The binding of both p53 mutants caused loss of intensity (peak broadening) to specific residues localized to a β-hairpin protruding from the ZFLR domain of Ydj1 (residues 168-187, which correspond to residues 168-184 in DNAJA2), with no interaction detected with CTDI or CTDII (Figures 3A and S4A-B).

To verify that this site is structure- and not sequence-specific, we repeated the binding assays to DNAJA2 homologue, this time using peptides that correspond to β4 and β9 DNAJA2-binding sites on p53. Addition of each of these peptides, that lack any secondary structure, to ^2^H, ^15^N-labeled DNAJA2 monomer, showed clear binding to the canonical CTDI and CTDII client binding sites on the chaperones, with no changes detected in the β-hairpin region (Figure S4C-D). Thus, while DNAJA2 CTDI and CTDII sites bind unfolded and misfolded clients ^46–48^, the β-hairpin instead recognizes structured β-sheet rich regions, most likely through re-stabilization of the hydrogen bonds.

To further elucidate the role of this new client-binding domain in class A JDPs, we generated a β-hairpin deletion mutant of DNAJA2 (DNAJA2^Δ169–182^, termed DNAJA2^ΔBH^) (Figure 3B). Removal of the β-hairpin completely abolished the ability of the human DNAJA2 chaperone to interact with the misfolded p53 mutants or prevent their aggregation *in vitro* (Figures 3C and S4E-F).

Thus, the DNAJA2 β-hairpin region represents a previously undiscovered client-binding fold in JDP chaperones, capable of recognizing proteins in their near-folded states at the very initial stages of misfolding.

### DNAJA2 ZFLR β-hairpin is essential for p53 interaction, but not for other chaperoning functions

We next tested whether the β-hairpin region also plays a crucial role in other DNAJA2 cellular functions. Deletion of this region had no effect on the ability of DNAJA2 to prevent the amyloid formation of the Alzheimer’s-associated protein tau (Figure S4G) and similar aggregation-prevention activity was observed for the wild-type and mutant DNAJA2s. This is in agreement with the recent finding that the chaperone interaction with tau monomers and fibers is mediated through CTDI and II domains ^48^. The DNAJA2^ΔBH^ mutant likewise remained as efficient as WT DNAJA2 in stimulating Hsp70 ATP hydrolysis rates (Figure S4H).

We also tested the ability of the mutant DNAJA2 to prevent the aggregation of insulin, whose structure contains only α-helical elements, and lacks the β-strand folds recognized by the chaperone in oncogenic p53 mutants. The DNAJA2^ΔBH^ mutant was indeed as efficient as the WT DNAJA2 chaperone in preventing insulin aggregation (Figure 3D), further indicating that class A JDP β-hairpin regions are specifically involved in the aggregation-prevention of β-strand rich proteins.

Lastly, we tested the ability of DNAJA2^ΔBH^ to support refolding of misfolded proteins as part of the Hsp70 chaperone system. The reactivation of chemically-denatured firefly luciferase (FFL) by Hsp70, Hsp110, and either DNAJA2 or DNAJA2^ΔBH^ was monitored by luminescence measurements upon addition of luciferin (Figure S4I). The Hsp70 system, working with WT DNAJA2 yielded 79±5% functional FFL, in agreement with previous reports ^51^. Interestingly, refolding activity of the system was only slightly reduced by the introduction of DNAJA2^ΔBH^ mutant (65±9%), demonstrating that the β-hairpin region of the chaperone does not play a significant role in FFL refolding (Figure S4I).

Thus, the DNAJA2 β-hairpin represents a novel functional element in class A JDPs, essential for recognizing initial misfolding, and preventing the aggregation of β-sheet rich proteins, such as p53.

To confirm this key role of the β-hairpin region in cells, we tested the interaction of overexpressed p53 R249S by co-immunoprecipitation with either WT DNAJA2 or mutant DNAJA2 lacking the β-hairpin. p53 R249S was found to co-precipitate only with WT DNAJA2 and no interaction was detected for DNAJA2^ΔBH^ (Figure 3E). Super-resolution confocal microscopy further showed that DNAJA2 colocalizes with residual p53 R249S aggregates, while no such colocalization in fluorescence signal was detected between p53 aggregates and DNAJA2^ΔBH^ (Figure S5A). The β-hairpin region is thus the main driver of interactions between DNAJA2 and mutant p53 in cells.

Accordingly, deletion of the β-hairpin also abolished the aggregation-prevention activity of DNAJA2 in cells, with cytosolic mutant p53 foci observed at the same frequency as the empty vector control (∼75%) (Figures 3F and S5B). This in contrast to the strong reduction in foci upon overexpression of WT DNAJA2 (49% and 45% for R249S and R282W respectively) despite equivalent over-expression levels of both p53 mutants and DNAJA2 variants under all conditions tested (Figure S5C). Fractionation of the cell lysate into soluble and insoluble fractions confirmed this observation, with the amount of mutant p53 found in the insoluble pellet fraction not significantly different in the presence of DNAJA2^ΔBH^ compared to the control (Figure S5D).

Combined, our results establish that it is the DNAJA2 ZFLR β-hairpin that specifically interacts with destabilized p53 mutants (and potentially other, similarly misfolded β-strand containing substrates), preventing their misfolding and subsequent aggregation, both *in vitro* and in human cell lines. Furthermore, while deletion of this region abolishes DNAJA2 chaperoning activity towards mutant p53, it only minimally affects other known functions of the chaperone.

### DNAJA2 sequesters non-native p53 into oligomeric complexes

Class A JDPs assemble into functional dimers in solution (Figure 3A), with each protomer containing a β-hairpin client binding site. As the presence of multiple client binding units was shown to greatly increase the affinity of many JDPs to their clients ^47, 52, 53^ the same could also be true for DNAJA2-p53 interaction.

To test whether two functional β-hairpin units are required for the DNAJA2–p53 interaction and aggregation-prevention function, we engineered a dimerization-deficient mutant (DNAJA2^mono^) by removal of the C-terminal dimerization domain ^54^ (Figure 3B, bottom). Preventing dimerization reduced DNAJA2 affinity for the destabilized p53 R282W mutant ∼5-fold (Figure 3G), and the monomeric chaperone showed no aggregation prevention activity towards these variants (Figure 3H). Thus, DNAJA2 prevention of the initial misfolding of β-sheet rich client proteins appears to require the simultaneous presence of both β-hairpin binding domains in the dimeric chaperone.

Structural information for the dimeric DNAJA2 in complex with p53 mutants, however, could not be obtained from the NMR experiments, due to severe peak broadening that was evident ∼30 minutes into the experiment. As such broadening is usually indicative of very high molecular weights, we used size-exclusion chromatography to learn more about the size of the p53-DNAJA2 assemblies. Surprisingly, the p53-DNAJA2 complexes eluted in much earlier fractions than would be expected from the combined molecular weights of p53 mutant DBD (23 kDa) and the DNAJA2 dimer (92 kDa) - indicating the formation of large oligomers (Figures S6A-B). No such behavior was detected for DNAJA2 alone, which remains dimeric even at high concentration and following a 5-hour incubation period at 37°C (Figure S6A, red dashed and solid lines). Multi-angle light scattering coupled with SEC (SEC-MALS) analysis showed these particles to be 1.6-2.3 MDa in molecular weight, with the DNAJA2-p53 assembles remaining stable even upon a second separation on SEC (Figure S6C). Dynamic light scattering (DLS) measurements of the complexes showed monodispersed particles with a hydrodynamic radius of 23nm (Figure S6D). Similar monodisperse particles were also detected with higher DNAJA2:p53 ratios (1-5 fold excess of DNAJA2), in line with these being stable oligomeric assemblies, rather than aggregates (Figures S6E-F).

Negative-stain EM images further confirmed that R249S-DNAJA2 and R282W–DNAJA2 form large structures with well-resolved particles, ∼40 nm in diameter, which are in stark contrast to the large amorphous aggregates formed by R249S and R282W p53 mutants in the absence of DNAJA2 chaperones (Figures S6G-J). It is worth noting that such a phenomenon is common to other chaperone systems, such as small heat shock proteins (sHSPs). The structural plasticity and increased client-binding affinity associated with the presence of multiple client-binding domains may thus be a common mechanism employed by non-ATP-dependent chaperones to prevent improper protein folding and aggregation^55, 56^.

### Hsp70 machinery disassembles the p53-DNAJA2 complexes releasing monomeric p53

Our data leads to the conclusion that DNAJA2 chaperones, despite being stable dimers in solution, form large oligomeric complexes upon interaction with non-natively folded p53. These complexes remain stable upon separation on size exclusion chromatography (SEC), with no visible release of p53, either as monomers or aggregates during the run (Figures S6B-C). As the DNAJA2 complexes do not disassemble spontaneously, we hypothesized that client-protein release must be dependent on downstream chaperone machineries. Hsp70s are the most likely candidates, as these chaperones were recently reported to disassemble highly ordered DNAJA2 tubular structures ^57^, and are known to directly interact with the J-domain of DNAJA2 ^50, 51, 58^.

To test the effect of the Hsp70 machinery on the p53-DNAJA2 complexes, preassembled R249S or R282W p53 DBD-DNAJA2 complexes were incubated with the different components of the Hsp70 system, along with a constant supply of ATP. The products of this incubation were then separated into soluble and insoluble fractions, with the soluble fraction further subjected to SEC, and blotted to detect the release of monomeric p53 proteins (Figure 4A). Control experiments established that, as expected, the mutant p53 DBD, incubated without chaperones, formed insoluble aggregates (Figures 4B-C; row 1), while R249S-DNAJA2 and R282W-DNAJA2 assemblies, incubated without the Hsp70 system, eluted as ∼1MDa complexes with no spontaneous release of monomeric p53 DBD detected in the low molecular weight fractions (Figures 4B-C; row 2).

**Figure. 4.**
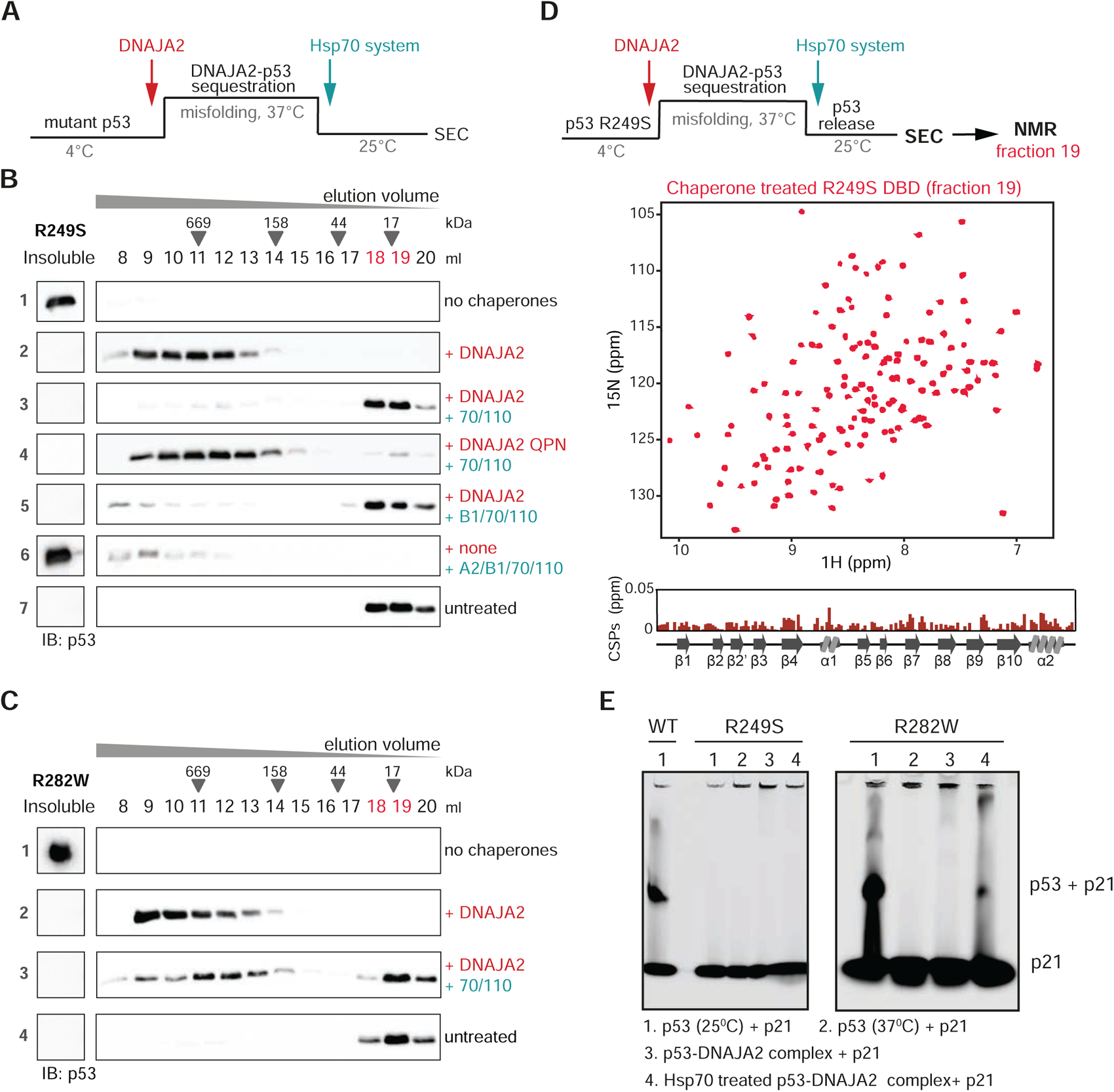
Release of p53 monomers from the p53-DNAJA2 complex. **(A-C)** Effect of various components of the Hsp70 chaperone system on the release of monomeric p53 mutants from pre-formed DNAJA2-p53 sequestration complexes. Misfolded p53 R249S **(B)** or R282W **(C)** mutants were incubated for 3h at 37°C either alone (row 1), in the presence of DNAJA2 WT (row 2) or DNAJA2 QPN (row 4). The formed DNAJA2-p53 complexes were then incubated with the indicated components of the Hsp70 system (rows 3-6), and the reactions were allowed to proceed at 25°C. The final products of these reactions were separated into insoluble and soluble fractions, with the soluble fraction further separated by SEC. All of the fractions were blotted against p53. The elution volumes of molecular standards are indicated above and the monomeric elution volume of untreated p53 (row 7) is marked in red. Addition of Hsp70 refolding system to p53-DNAJA2 complexes is sufficient to efficiently release soluble p53 monomers. **(D, top)** ^1^H - ^15^N HSQC NMR spectra of ^15^N-labeled p53 R249S DBD following sequestration by DNAJA2 and release by the Hsp70-system (fractions 18-19 in row 3 of panel B). **(D, bottom)** Difference in chemical shifts between the untreated p53 R249S DBD and the DNAJA2/Hsp70/Hsp110 treated protein. No significant chemical shift perturbations are detected between the two samples indicating that the chaperones do not affect the conformation of the released p53 mutant. **(E)** Interaction of p53 WT, R249S, and R282W proteins with p21 promoter DNA as detected by EMSA. p53 variants were incubated at elevated temperatures (37°C) alone and in the presence of DNAJA2 chaperones. Following incubation, Hsp70 system was added to release the monomeric p53. The untreated p53 incubated at 25°C was used as a control. Both the WT-untreated p53 and the chaperone-treated protein bind to DNA, while the aggregated protein and p53 in complex with DNAJA2 do not. The position of the p53-p21 complex and unbound p21 DNA are indicated. See also Figures S6.

Addition of the Hsp70 system (Hsp70 and the nucleotide exchange factor Hsp110), to samples containing R249S or R282W p53-DNAJA2 complexes, resulted in a strong enrichment of the monomeric p53 DBD population, with over 90% of the total p53 DBD eluting in the low molecular weight fractions (Figures 4B-C; row 3). This complex-disassembly required a direct interaction between Hsp70 and the DNAJA2 J-domain, as mutation to this region (DNAJA2^QPN^) abolished the Hsp70-dependent release activity (Figure 4B; row 4).

Further experiments identified that the levels of the released monomeric p53 were not increased (∼85% p53 monomers) upon addition of the mixed-JDP Hsp70 disaggregation machinery (Hsp70, DNAJA2, DNAJB1, and Hsp110) ^59^ (Figure 4B; row 5) nor could this machinery disaggregate p53 aggregates preformed in the absence of DNAJA2 (Figure 4B; row 6), confirming that p53-DNAJA2 complexes are oligomeric assemblies, rather than soluble aggregates requiring disaggregation.

To determine the conformation of the client protein following its release from DNAJA2 sequestration, we subjected SEC fractions containing released ^15^N-labeled monomeric p53 R249S (Figure 4B, row 3, fractions 18-19) to NMR measurements. The ^1^H-^15^N HSQC spectrum of the chaperone-treated R249S p53 was characteristic of a properly folded protein (Figure 4D, top) and highly similar to that of the untreated p53 R249S DBD mutant (Figure 4D, bottom). Since, in the absence of Hsp90 chaperones, the Hsp70 system is not sufficient to refold unfolded p53 ^25, 26^ to a mature and functional protein, our evidence for a properly folded p53 DBD suggests that the DNAJA2 chaperones sequester the destabilized p53 in a folded or nearly-folded conformation primed for release by Hsp70.

### DNAJA2 stabilizes the oncogenic conformation of p53 mutants, protecting them from degradation

Our findings describe a new mode of function for class A JDPs, which recognize their misfolded clients through the detection of increased dynamics and transient breakage of hydrogen bonds in β-sheet rich proteins. Oligomeric DNAJA2 structures then sequester the misfolded proteins, protecting them potentially until stress conditions are relieved and Hsp70 chaperones become available for disassembly of the DNAJA2-substrate complex. These chaperoning functions are most likely beneficial for the cell and ensure protein homeostasis and cell survival under various stress conditions. However, in the case of mutant p53 proteins this DNAJA2 activity may in fact be detrimental to the host.

In cancer cells, molecular chaperones increase the stability and protect the oncogenic p53 mutants from proteasomal degradation ^12, 60–62^. This results in accumulation of high cellular levels of mutant p53, crucial for its oncogenic gain of function (GOF) activities ^16,17, 61, 63^. Interestingly, such a tumor-protective function was recently described for the DNAJA1 chaperone, which is a homologue of DNAJA2 bearing 67% sequence similarity. DNAJA1 depletion enhanced the degradation of mutant p53 and substantially reduced the malignant properties of cancer cells ^64–67^. In light of these findings, we hypothesized that our newly identified interaction of DNAJA2 chaperones with mutant p53 variants may also stabilize and protect the mutant forms of the protein from proteasomal degradation, thus promoting its GOF cancer-associated effects ^3^.

To test this, we first confirmed that DNAJA2 chaperones stabilize the non-functional, oncogenic form of the mutant p53s. Indeed, the R249S p53 mutant released from DNAJA2 sequestration did not regain any DNA-binding activity, as evident from electrophoretic mobility shift assays (EMSA) with Cy5.5-labeled double-stranded p21 DNA (Figure 4E), which contains native p53 binding sites ^68, 69^. The DNA-binding properties of chaperone-treated R282W mutant and heat-destabilized WT p53 were likewise found to be similar to the untreated proteins (Figure 4E). Furthermore, overexpression of DNAJA2 in SaOS-2 cells expressing the p53 mutants, while reducing p53 aggregation, did not induce the expression of p53 endogenous target genes (p21 and MDM2), as monitored by qPCR (Figures S7A-B). Thus, DNAJA2 binding stabilizes the oncogenic conformations of p53 (rather than facilitating refolding to native-like p53), protecting the proteins from aggregation while potentially preserving the GOF effects of these mutants.

To determine whether sequestration by DNAJA2 chaperones protects mutant p53 from degradation, we quantified the levels of endogenously expressed R282W p53 mutant in pancreatic cancer cell lines upon knockdown of class A JDPs and overexpression of either WT DNAJA2 chaperone or DNAJA2^ΔBH^, lacking the p53-binding β-hairpin region. As expected, knockdown of class A JDPs reduced the levels of R282W p53 (Figures S7C-E), and substantially enhanced the rates of p53 degradation in cycloheximide chase (CXH) experiments (Figures 5A-B). These rates, however, significantly decreased upon overexpression of DNAJA2, suggesting that this chaperone indeed protects p53 from proteasomal degradation (Figures 5A-B and S7E). Importantly, overexpression of DNAJA2^ΔBH^, lacking the β-hairpin region, did not lead to a decrease in the degradation rates of R282W p53 (Figures 5A-B), and the overall steady-state protein levels of the mutant were significantly reduced (Figure S7D). p53 degradation was completely inhibited upon overexpression of DNAJA2^QPN^ carrying mutations in the conserved HPD region of the J-domain (Figures 5A-B), which is essential for Hsp70 recruitment and activation, indicating that the DNAJA2-Hsp70 interaction is required for the release of p53 and its subsequent delivery to the proteasome.

**Figure 5.**
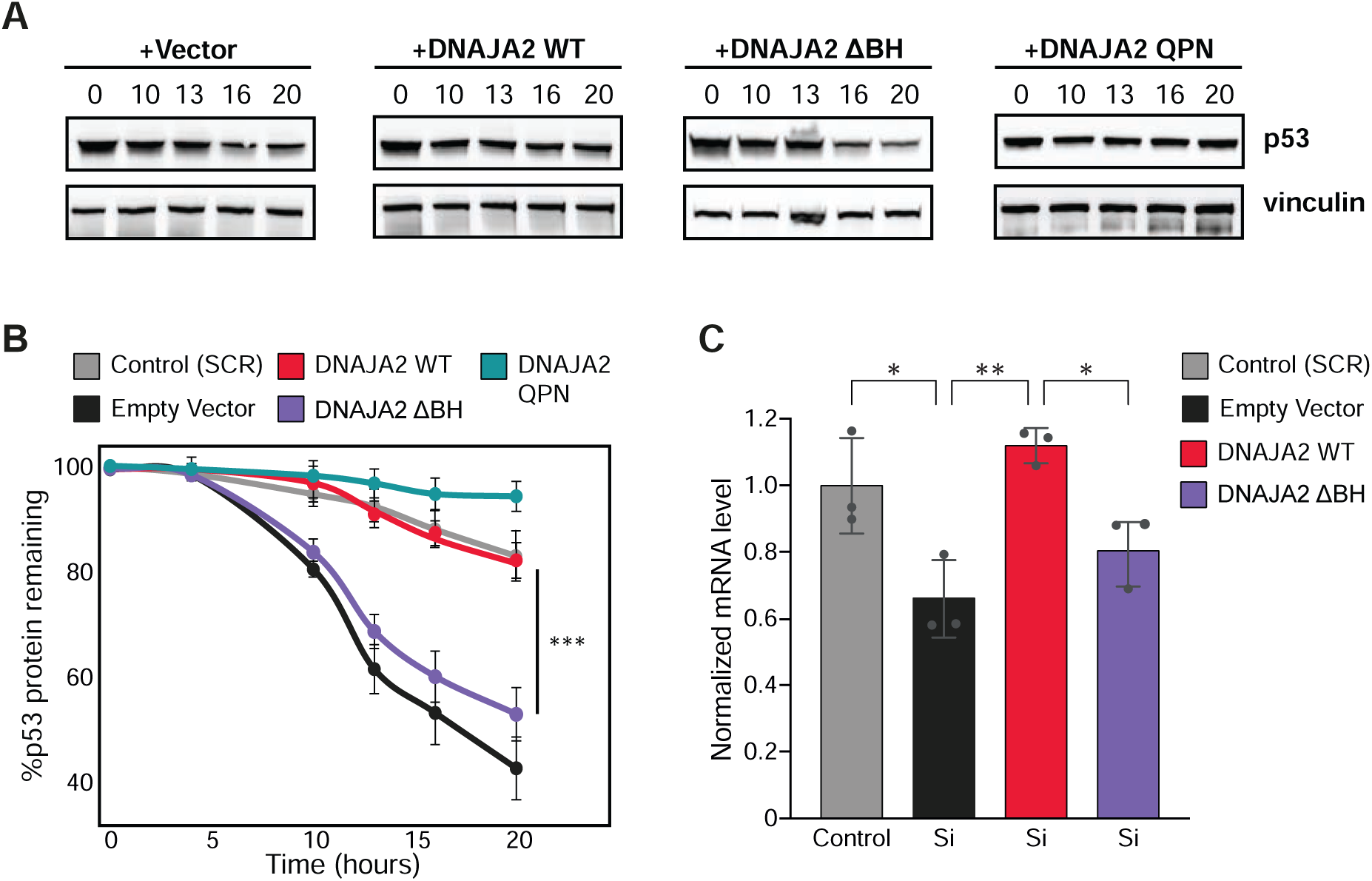
Class A JDP chaperones protect the oncogenic p53 mutants from degradation. **(A)** Cycloheximide (CHX) chase assay to monitor the degradation of endogenous p53 mutant (R282W) in PaTu 8988 cells following depletion of class A JDPs (siDNAJA1) and overexpression of DNAJA2^WT^, DNAJA2^ΔBH^, or DNAJA2^QPN^ chaperones. Cells were immunoblotted for p53 at the indicated time points following treatment with CHX (0-20 hours). Vinculin was monitored as a control. **(B)** Quantification of p53 levels in panel A. The graph shows the ratio of the relative levels of p53 and vinculin (control) at each time point; time 0 was set to 100%. Data represent mean values ± s.d (n=3). *** p<0.001 (one-way ANOVA test). See also Figure S7. **(C)** PLK2 mRNA levels quantified 60 h post-treatment and normalized to GAPDH levels. Data represent mean values ± s.d (n=3). * p<0.05, ** p<0.01 (one-way ANOVA test).

Combined, these results demonstrate that the interaction of class A JDPs with mutant p53, carried out through their unique β-hairpin region, both stabilizes the protein and protects it from proteasomal degradation (Figure 6A).

**Figure 6.**
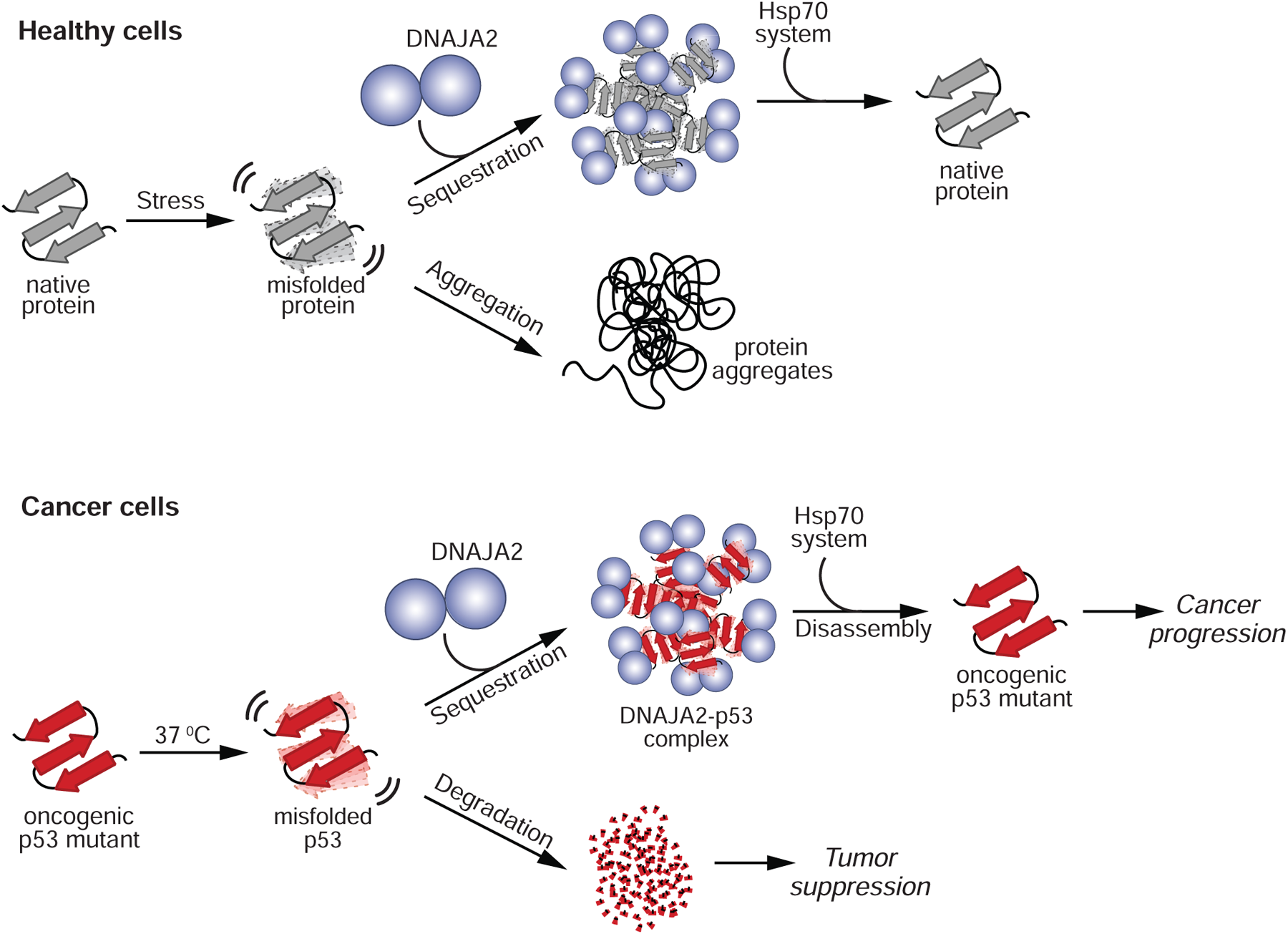
Model of class JDP function in health and disease. **(A)** Cartoon model for class A JDP chaperone function in both healthy and cancer cells (top) Class A JDPs, such as DNAJA2, can recognize initial stages of misfolding in β-sheet rich proteins, via their β-hairpin client binding site. Once bound to the misfolded proteins, the chaperones assemble into large oligomeric complexes, sequestering their clients and protecting them from aggregation. After the cellular stress has abated, the Hsp70 chaperone system disassembles the chaperone-client complexes, releasing functional native proteins. (bottom) In cancer, however, class A JDPs also bind to the destabilized, oncogenic mutants of p53, protecting these oncoproteins, through sequestration, from degradation by cellular protein quality control machineries. This action enables the continued function of the oncogenic p53 mutants, effectively promoting the progression of cancer.

Next, we tested if eliminating the mutant p53 interaction and stabilization by DNAJA2 chaperones, thus enhancing its degradation, will reduce the p53 mutant GOF ability to facilitate the expression of other oncogenic proteins. To this end we followed the transcription levels of an oncogenic protein, Polo-like kinase 2 (PLK2) ^70, 71^ a known target gene of R282W mutant p53 in malignant cells^72^. As expected, PLK2 mRNA levels were significantly reduced following the knockdown of class A JDPs (Figure 5C). Overexpression of DNAJA2 WT in PaTu cells fully rescued the mRNA levels of PLK2, while similar expression of but not DNAJA2^ΔBH^ mutant, unable to bind and stabilize the mutant p53, had no effect (Figure 5C). Thus, DNAJA2 chaperones, via their β-hairpin domain, bind and stabilize the mutant p53, protecting it from degradation and thereby promoting the oncogenic properties of mutant p53-carrying cancers.

## Discussion

JDP chaperones, together with their Hsp70 partners, play crucial roles in maintaining proteostasis in the cell, protecting against protein misfolding induced by stress, disease, and aging. Little, however, was known about how J-domain chaperones interact with and affect their clients, which are primarily destabilized, misfolding-prone proteins.

Conformational p53 oncogenic mutants are misfolding-prone clients of the chaperone system. Specifically, several JDP family members were described to promote cancer by acting on these destabilized mutant proteins, preventing p53 ubiquitination and degradation ^3, 13, 14^. Here we identified a previously uncharacterized client binding domain in class A JDPs, which can recognize the initial stages of misfolding in p53 mutants and, via direct interaction, stabilize the β-sheet rich DNA-binding domain in these oncoproteins. Through this chaperone site, consisting of two β-hairpins, class A JDPs can sense the increase in dynamics and transient breakage of hydrogen bonds in the mutant protein and stabilize it before it misfolds. Such a mode of recognition represents an entirely novel functional mechanism for molecular chaperones, which to date have only been shown to recognize hydrophobic regions exposed following large structural rearrangements in already highly destabilized or unfolded clients ^1,39^. Moreover, as it is decidedly more difficult to refold β-sheet proteins once they have misfolded, we propose that this early detection mechanism, may be widely employed by this subfamily to protect such β-sheet rich proteins in the cell. As exposure of β-sheet regions is known to induce protein aggregation, and is strongly associated with many protein misfolding and conformational diseases ^73, 74^, class A chaperones may thus form the cell’s first line of defense during stress.

It is important to note that class A JDPs also contain additional client binding domains located in their CTDs (CTDI and CTDII), that are known to bind unfolded polypeptides enriched in hydrophobic and aromatic residues ^46, 47^, and are essential for folding and refolding of largely unstructured proteins, as part of the Hsp70 system. In addition, the CTDs have been reported to play a role in the aggregation-prevention activity of class A JDPs, by suppressing toxic amyloid aggregation, such as in the case of the Alzheimer’s-associated protein, tau ^48, 75^. Unlike the CTDs, the β-hairpin site is not required for interaction with unfolded proteins, protein refolding, or amyloid aggregation prevention. The β-hairpin is, however, the only site capable of sensing the increase in dynamics and stabilizing β-sheet rich proteins before they misfold and aggregate. It is therefore unsurprising that class A JDPs, being the only chaperones with such a domain, are likewise the only J-domain proteins which can prevent the misfolding of destabilized p53 mutants. Overall, it would appear that it is this combined action of the different client-binding sites in class A JDPs that confer their ability to bind and chaperone a broad range of protein conformers in the cell.

We have also uncovered that, while the class A DNAJA2 normally forms dimers, this chaperone assembles into large oligomeric complexes upon interaction of the β-hairpin region with misfolded clients. These sequestration complexes remain both stable and soluble, and are vital to protecting the trapped clients from further misfolding and aggregation. Such a mode of function is reminiscent of small heat shock proteins ^56^, and no such sequestration assemblies have previously been described for any other proteins in the Hsp70 chaperone system. We therefore propose that this class-A JDP sequestration mechanism ^57^, is widely used by cells to prevent misfolding and aggregation of β-sheet rich proteins during extreme conditions. After the cellular stress has abated, the protective DNAJA2-p53 complexes are then disassembled by the Hsp70 system, releasing folded monomeric p53 (Figure 6A). Interestingly, we find that the client proteins are kept in a nearly native folded state once in complex with the DNAJA2 chaperones and are thus released as native monomeric proteins without needing to be refolded by the Hsp70/Hsp90 chaperone systems. Furthermore, this mechanism may be of particular importance under stress conditions, where Hsp70 levels are limited, and other protective mechanisms are required to prevent protein aggregation.

This same mechanism, however, poses a significant risk in the case of mutant p53, where these protective functions of the JDP chaperones prevent its clearance by degradation and thus can support the anti-apoptotic, tumor-promoting properties of the oncogenic variants (Figure 6A). These findings highlight the double-edged nature of chaperoning activities in the context of disease, where normally protective functions can be disruptive to health if not coupled to effective clearance of the faulty client proteins. Based on these observations, we envisage that inhibitors specifically targeting the β-hairpin region in class A JDPs could reduce the levels of destabilized oncogenic p53 mutants, while not affecting other broad housekeeping chaperoning functions in the cell. It is important to note that while JDP-based cancer therapeutics were already suggested in the literature ^66, 76–78^, such approaches target the J-domain region in the JDPs, thus simultaneously affecting the interaction of all ∼50 J-domain proteins with Hsp70, potentially leading to severe side effects. In contrast, the inhibition of the β-hairpin represents a novel target with the possibility of significantly improved outcomes, as it focuses on a site present in only a small subset of JDPs (class A), and participates specifically in the stabilization of β-sheet rich proteins, such as p53. Such an approach could potentially restrict the malignant properties of cancer cells, while presenting minimal side effects.

## SUPPLEMENTAL INFORMATION

Supplemental information includes seven figures.

## ACKNOWLEDGEMENTS

The authors would like to thank C.S. Tran for cell culture and microscopy assistance, T. Scherf for NMR support and the Clore Institute for High-Field Magnetic Resonance Imaging and Spectroscopy. We also thank V. Rotter, M. Oren, Z. Shakked, J. Tittelmeier, C. Nussbaum-Krammer, M.P. Latham, and D.F. Hansen for helpful discussions and advice, and N. Nillegoda for contributions to funding acquisition. R.R. is supported by the European Research Council starting grant (ERC-2018-STG 802001), the Minerva Foundation, and a research grant from the Blythe Brenden-Mann New Scientist Fund. R.R. and B.B are supported by the Germany - Israel Cooperation - IMOS - DKFZ - Cooperation in Cancer Research. BB is supported by grant BU617/21-1 from the Deutsche Forschungsgemeinschaft.

## AUTHOR CONTRIBUTIONS

G.Z., L.D., A.W., B.B., and R.R. designed the research; G.Z., M.S., and R.R. performed the NMR spectroscopy measurements, processed and analyzed the data. G.Z. and O.F., performed the biochemical and functional assays; L.D. preformed all the cellular aggregation assays and imaging. M.K., G.Z., and T.I. carried out the cellular degradation assays. G.Z., L.D., M.K., O.F., A.W., and R.R. analyzed data; and G.Z., L.D., O.F., A.W., B.B., and R.R. wrote the paper.

## DECLARATION OF INTERESTS

The authors declare no competing interests.

### Data and material availability

All data are presented in the paper and/or the supplementary materials. NMR chemical shifts for human p53 DBD have been deposited in the Biological Magnetic Resonance Data Bank accession code 51753.

