## Supplemental methods and figures for "A unique chaperoning mechanism in Class A JDPs recognizes and stabilizes mutant p53"

#### Key resources table

| REAGENT or RESOURCE | SOURCE | IDENTIFIER |
|---|---|---|
| **Antibodies** | | |
| Mouse monoclonal (PAb 240) anti-p53 | abcam | Cat#ab26; RRID:AB_303198 |
| Mouse monoclonal (DO-1) anti-p53 | Santa Cruz | Cat#sc-126; RRID:AB_628082 |
| Rabbit monoclonal (EPR8185) anti-Vinculin | abcam | Cat#ab129002; RRID:AB_11144129 |
| Rabbit polyclonal anti-DNAJA1 | abcam | Cat#ab192904 |
| Rabbit monoclonal (SV5-P-K) anti-V5 tag | abcam | Cat#ab206566; RRID:AB_2819156 |
| Rabbit monoclonal (DrH8Q) anti-V5 tag | Cell Signaling | Cat#13202; RRID:AB_2687461 |
| Mouse monoclonal (E10) anti-V5 tag | Santa Cruz | Cat#sc-81594; RRID: AB_1131162 |
| Rabbit monoclonal (60A8) anti-c-Jun | Cell Signaling | Cat#9165; RRID: AB_2130165 |
| Mouse monoclonal anti-Actin | Invitrogen | Cat#MA5-11869; RRID: AB_11004139 |
| Rabbit polyclonal anti-DNAJA2 | Invitrogen | Cat#PA5-65583; RRID:AB_2662184 |
| Goat anti mouse IgG-HRP | JIR | Cat#115-035-003; RRID: AB_10015289 |
| Goat anti rabbit IgG-HRP | JIR | Cat#111-035-003; RRID: AB_2313567 |
| Goat anti-mouse Alexa 568 | Invitrogen | Cat#A-11004; RRID: AB_2534072 |
| Goat anti-rabbit Alexa 488 | Invitrogen | Cat#A32731; RRID: AB_2633280 |
| Goat anti-rabbit Alexa Fluor488 | abcam | Cat#ab150077; RRID:AB_2630356 |
| **Bacterial and virus strains** | | |
| BL21(DE3) Competent Cells | Novagen | Cat#69450 |
| Rosetta™(DE3) Competent Cells | Novagen | Cat# 70954 |
| **Chemicals, peptides, and recombinant proteins** | | |
| Thioflavin T | Thermo Scientific | Cat#211760050; CAS: 2390-54-7 |
| Luciferase assay system | Promega | Cat# E1500 |
| MG-132 | Cell Signaling | Cat#21945S; CAS: 133407-82-6 |
| Cycloheximide | Sigma-Aldrich | Cat#C7698; CAS: 66-81-9 |
| DAPI | Sigma-Aldrich | CAS: 28718-90-3 |
| DAPI | Invitrogen | Cat#D1306; CAS: 28718-91-4 |

| | | |
|---|---|---|
| Lipofectamine 3000 | Invitrogen | Cat#L3000001 |
| cOmplete Protease Inhibitor Cocktail | Roche | Cat#11697498001 |
| MDCC | Adipogen | Cat#CDX-D0198;<br>CAS: 156571-46-9 |
| Insulin | Sigma-Aldrich | Cat#91077C;<br>GenPept:P01308 |
| Heparin | Sigma-Aldrich | Cat#H3393;<br>CAS: 9041-08-1 |
| Scrambled RNA | Dharmacon | Cat# D-001810-10-05 |
| Penicillin-Streptomycin Solution | Biological Industries | 03-031-1B |
| Alexa Fluor 488 C5 Maleimide | Invitrogen | Cat# A10254 |
| DMEM | Gibco | Cat#41965-039 |
| ExpressPlus PAGE | GenScript | Cat#M42012 |
| NativePAGE Bis-Tris | Invitrogen | Cat#BN1001BOX |
| ProLong Gold Antifade | Invitrogen | Cat#P36930 |
| $^{15}NH_4Cl$ | CIL | Cat#NLM-467 |
| $^{13}$C-D-glucose | CIL | Cat#CDLM-3813 |
| D-glucose | Sigma-Aldrich | Cat#552003 |
| Deuterium oxide | Sigma-Aldrich | Cat#151882 |
| Creatine kinase | Sigma-Aldrich | Cat#C7886 |
| 4-20% gradient SDS-PAGE | GenScript | Cat#M00656 |
| ECL | Bio-Rad | Cat#1705060 |
| Dynabeads protein G | Invitrogen | Cat#10003D |
| EM grids | Electron Microscopy Sciences | Cat#CF300-CU |
| Uranyl-acetate | | |
| Gel filtration standards | Bio-Rad | Cat#1511901 |
| p53 β4 peptide (residues 155-164)<br>TRVRAMAIYK | GenScript | Custom made |
| p53 β9 peptide (residues 248-263)<br>RSPILTIITLEDSSGN | GenScript | Custom made |
| **Critical commercial assays** | | |
| RNeasy Mini Kit | Qiagen | Cat#74104 |
| iScript cDNA Synthesis Kit | Bio-Rad | Cat#1708890 |
| TB Green Premix Ex Taq II | Takara | Cat#RR82WR |
| Wizard Plus SV Minipreps DNA Purification System | Promega | Cat#A1460 |
| qScript cDNA Synthesis Kit | Quantabio | Cat#95047 |
| Fast SYBR™ Green Master Mix | Applied Biosystems | CAT#4385612 |
| QIAGEN Plasmid Maxi Kit | Qiagen | Cat#12163 |
| Pierce™ BCA Protein Assay Kit | Thermo Scientific | Cat#23225 |
| **Deposited data** | | |
| p53 DBD (94-293) NMR backbone assignments | This paper | BMRB: 51753 |

| | | |
|---|---|---|
| **Experimental models: Cell lines** | | |
| PaTu 8988t | | N/A |
| SaOS-2 | ATCC | ATCC - HTB-85; RRID:CVCL_0548 |
| **Oligonucleotides** | | |
| Cy5.5 labeled DNA-oligo p21 TGGCCATCAGGAACATGACCCAACATGTTGA GCTCTGGCA | Sigma-Aldrich | N/A |
| siDNAJA1 | Dharmacon | Cat#L-019617 |
| p21 primers for qPCR<br>forward CGCTAATGGCGGGCTG<br>reverse CGGTGACAAAGTCGAAGTTCC | This paper | N/A |
| MDM2 primers for qPCR<br>forward ACCTCACAGATTCCAGCTTCG<br>reverse TTTCATAGTATAAGTGTCTTTTT | This paper | N/A |
| PLK2 primers for qPCR<br>forward AAGTTGGGGACTTCGGTCTGG<br>reverse TGGCAGGAGCCAGCAATGA | This paper | N/A |
| ACTB primers for qPCR<br>forward CCTTTGCTTGCTTTCTTTCC<br>reverse AGAGAAGTGGGGTGGCTTTT | This paper | N/A |
| GAPDH primers for qPCR<br>forward GGTCGGAGTCAACGGATTTGG<br>reverse ACTCCACGACGTACTCAGCG | This paper | N/A |
| **Recombinant DNA** | | |
| pET22b PBP (A197C) | [82] | Addgene 78198 |
| pET29b DNAJA2 TEV cleavable His tag | [58] | N/A |
| pET29b DNAJA2$^{QPN}$ H34Q, D36N TEV cleavable His tag | This paper | N/A |
| pET29b DNAJA2$^{\Delta BH}$ Δ169-182 TEV cleavable His tag | This paper | N/A |
| pET29b DNAJA2$^{mono}$ (111-353) F337D TEV cleavable His tag | This paper | N/A |
| pcDNA5 DNAJA2 with V5 tag | [83] | Addgene 19519 |
| pcDNA5 DNAJA2 $^{\Delta BH}$ Δ169-182 with V5 tag | This paper | N/A |
| pET29b DNAJA1 TEV cleavable His tag | This paper | N/A |
| pET-sumo DNAJB1 His-SUMO tag Ulp1 cleavable | [59] | N/A |
| pcDNA5 DNAJB1 with V5 tag | [83] | Addgene 19522 |
| pET29b DNAJB2 His-SUMO tag Ulp1 cleavable | This paper | N/A |
| pET29b DNAJB4 TEV cleavable His tag | [58] | N/A |
| pET-sumo DNAJB6 His-SUMO tag Ulp1 cleavable | This paper | N/A |
| pET 29b DNAJC7 TEV cleavable His tag | [84] | N/A |
| pET29b DNAJC8 His-SUMO tag Ulp1 cleavable | This paper | N/A |
| pET-sumo Hsp70 His-SUMO tag Ulp1 cleavable | [52] | N/A |
| K151 Hsp110 His-SUMO tag Ulp1 cleavable | [59] | N/A |

| | | |
|---|---|---|
| pET-27b p53 DBD (94-293) | [85] | N/A |
| pcDNA3.1 p53 | [86] | Addgene 69003 |
| pET-27b p53 R249S DBD (94-293) | [42] | N/A |
| pcDNA3.1 p53 R249S | This paper | N/A |
| pET-27b p53 R282W DBD (94-293) | [44] | N/A |
| pcDNA3.1 p53 R282W | This paper | N/A |
| pET-Sumo tau C291S, C322S, L243C, T373C | [48] | N/A |
| pET SUMO Ydj1 mono (111-351) F335D His-SUMO tag Ulp1 cleavable | This paper | N/A |
| Ulp1 protease | produced in house | N/A |
| Tobacco Etch Virus Protease | produced in house | N/A |
| **Software and algorithms** | | |
| Image Studio | LI-COR | https://www.licor.com/bio/image-studio/resources#is5-clx |
| ImageJ | | https://imagej.nih.gov/ij/ |
| NMRFAM-SPARKY | [87] | https://nmrfam.wisc.edu/nmrfam-sparky-distribution/ |
| Topspin | Bruker | https://www.bruker.com/en/products-and-solutions/mr/nmr-software/topspin.html |
| NMRPipe | [88] | https://www.ibbr.umd.edu/nmrpipe/ |
| CcpNmrAnalysis | [89] | https://ccpn.ac.uk/software/version-2/ |
| Prism | GraphPad | https://www.graphpad.com/scientific-software/prism/ |
| Dynamics | Wyatt Technology | https://www.wyatt.com/products/software/dynamics.html |
| ASTRA | Wyatt Technology | https://www.wyatt.com/products/software/astra.html |
| UCSF ChimeraX | [90] | https://www.cgl.ucsf.edu/chimerax/download.html |

#### RESOURCE AVAILABILITY

#### MATERIALS AVAILABILITY

All unique/stable reagents generated in this study will be made available on request to the lead contact but may require a completed Materials Transfer Agreement.

### METHOD DETAILS

#### Construct Preparation

Codon optimized DNAJA2 wt and mutants, DNAJA1, DNAJB4, DNAJC7, and Hsp110 were cloned into pET-29b(+) vector with a N-terminal $His_6$ tag followed by a tobacco etch virus (TEV) protease cleavage site. DNAJB1, DNAJB2, DNAJB6, DNAJC8, Hsp70, and Ydj1 (residues 111-351, F335D; yeast orthologue of class A JDPs), were expressed from the pET-SUMO vector with an N-terminal $His_6$ purification tag. All mutations were introduced by QuickChange or TPCR.

Monomeric DNAJA2 ($DNAJA2^{mono}$, designed by homology to yeast Ydj1[46]) was generated by introducing a premature stop codon at position D353 and mutation of residue F337 to D337, by site-directed mutagenesis.

p53 DBD wt and mutants in pET27 vector were a generous gift from Zippora Shakked, Weizmann Institute of Science, Israel.

The mammalian expression plasmid pcDNA3.1-p53 was a gift from David Merk (Addgene plasmid #69003). The R249S and R282W mutations were introduced into the pcDNA3.1-p53 vector by PCR mutagenesis using Gibson Assembly cloning. Plasmids pcDNA5/FRT/TO V5-DNAJA2 (Addgene plasmid #19519) and pcDNA5/FRT/TO V5-DNAJB1 (Addgene plasmid #19522) were gifts from Harm Kampinga. pcDNA5 V5-$DNAJA2^{\Delta BH}$ was cloned by deleting the β-hairpin of pcDNA5 V5-DNAJA2 by PCR using Gibson Assembly cloning.

#### Protein expression and purification

p53 DBD (94-293) constructs were expressed in BL21(DE3) cells (Novagen) and grown in LB or M9-minimal media supplemented with $^{15}NH_4Cl$ at 37 °C until $OD_{600}$ of 0.6. Expression was induced by addition of 0.5 mM IPTG, supplemented with 100 µM $ZnCl_2$, and allowed to proceed overnight at 15 °C. Following harvesting, cells were lysed by French Press in 50 mM sodium phosphate buffer pH 7.1, 25 mM NaCl, and 5 mM DTT and purified on a 5 ml HiTrap SP column (GE Healthcare). The p53 DBD proteins were eluted in 50 mM sodium phosphate buffer (pH 7.1, with 5 mM DTT) using a 25-300 mM NaCl gradient over 150 ml. The proteins were further purified on a HiLoad 16/600 Superdex 75 pg size exclusion column (GE Healthcare). All proteins were stored at -80°C until use.

DNAJA1, DNAJB4, DNAJA2, $DNAJA2^{\Delta 169\text{-}182}$, and DNAJA2 monomer constructs were expressed in BL21(DE3) cells (Novagen) and grown in LB or M9-minimal $D_2O$ media supplemented with $^2H$-glucose grown at 37°C until OD 600 nm 0.8-1.0. Expression was induced by the addition of 1 mM of IPTG and allowed to proceed overnight at 25°C. Following harvesting, cells were lysed by French Press in 50 mM Tris, pH 8.0 with 750 mM KCl, 10 mM MgCl, 10% glycerol, 10 mM imidazole, 10 mM BME, and purified on a 5 mL HisTrap HP Ni-NTA column (GE Healthcare). The proteins were released from the column with 50 mM Tris, pH 8.0, 500 mM KCl, 250 mM imidazole, 10 mM $MgCl_2$ elution buffer. The 6His-tag was removed by an overnight cleavage with TEV protease at 4 °C. The cleaved proteins were further separated from the proteases and the uncleaved protein fractions by reverse capture Ni-NTA. The proteins were further purified on a HiLoad 16/600 Superdex 200 / 75 pg gel (GE Healthcare) filtration columns.

Human DNAJB1, DNAJB2, DNAJB6, DNAJC7, DNAJC8, Hsp70 (HSPA8), and s. cerevisiae $Ydj1^{111\text{-}351}$ constructs were expressed in *E. coli* BL21 as $His_6$-Smt3 (H6-sumo) fusion proteins. Proteins were purified by nickel affinity purification (Ni-NTA, GE-Healthcare), the tag was cleaved

by Ulp1 protease and the cleaved tag was removed by reverse nickel affinity purification. Proteins were further purified by size exclusion chromatography (HiLoad 16/600 Superdex 200, GE Healthcare). DNAJB4, DNAJC7, and Hsp110 were expressed in BL21 (DE3) and purified as previously described [52,58,84].
All proteins were concentrated, aliquoted, snap-frozen in liquid nitrogen and stored at −80 °C. The purity of all proteins was confirmed by SDS–PAGE and mass spec.

#### NMR Spectroscopy

All NMR experiments were carried out on 14.1T (600 MHz), 18.8T (800 MHz) or 23.5T (1000 MHz) Bruker spectrometers equipped with triple resonance single (z) or triple (x,y,z) gradient cryoprobes. The experiments were processed with Topspin 4.1 (Bruker) or NMRPipe [88] and analyzed with NMRFAM-SPARKY [87] and CcpNmrAnalysis [89].

Isotopically labeled proteins for NMR were grown in M9 $H_2O$ or $D_2O$ media supplemented with $^{15}NH_4Cl$ (and $^{13}C$-glucose) as the sole nitrogen (and carbon) source.

##### NMR Assignment experiments

Backbone $^1H$, $^{15}N$ and $^{13}C$ resonance assignments were carried out on a sample of 0.5 mM p53 DBD in 50 mM NaPi pH 7.2 buffer supplemented with 150 mM NaCl, 2.5 mM DTT, 0.02% $NaN_3$ and 10% $D_2O$ buffer. Assignments were obtained by recording HNCACB, HNCOCA, HNCA, HN(CA)CO, and HNCO experiments on an 800 MHz Bruker spectrometer. Unambiguous assignment of 94% of non-proline residues was achieved.

##### NMR binding experiments

p53 DBD interaction with chaperones was assayed for 200 μM samples of [U-$^{15}N$]-labeled p53 R249S or R282W in 50 mM NaPi pH 7.2, 75 mM NaCl, 2.5 mM DTT, 0.02% $NaN_3$, and 10% $D_2O$. The following concentrations of binding parents were used: 400 μM [U-$^1H$]-DNAJB1, 400 μM [U-$^1H$]-DNAJA1, 400 μM [U-$^1H$]-DNAJA2, 400 μM [U-$^2H$]-DNAJA2, or 400 μM [U-$^1H$]- DNAJA2$^{\Delta 169\text{-}182}$.

DNAJA2 interaction with p53 mutants was tested using a 150 μM sample of [U-$^2H$,$^{15}N$]-labeled Ydj1$^{111\text{-}351}$ and 300 μM [U-$^1H$]-p53 DBD variants in 50 mM HEPES pH 7.0, 60 mM NaCl, 1.0 mM DTT, 0.02% $NaN_3$, and 10% $D_2O$.

DNAJA2 interaction with β4 and β9 peptides of p53 were tested using a 150 μM sample of [U-$^2H$,$^{15}N$]-labeled Ydj1$^{111\text{-}351}$ and 800 μM of each [U-$^1H$]-peptide in 50 mM HEPES pH 7.0, 60 mM NaCl, 1.0 mM DTT, 0.02% $NaN_3$, and 10% $D_2O$.

$^1H$-$^{15}N$ HSQC-TROSY (transverse relaxation optimized spectroscopy) spectra were acquired for each sample and intensity ratios ($I/I_0$) were calculated, where I and $I_0$ correspond to the peak intensity of the bound and free samples, respectively. $I/I_0$ that were smaller than one standard deviation from the mean were considered significant.

##### CPMG relaxation dispersions

The $^{15}N$ CPMG relaxation dispersion data sets were recorded at 25 °C at $^1H$ frequency of 1000 MHz using a constant-time version of the relaxation-compensated TROSY CPMG pulse sequence [91] on the samples of 360 μM [$^2H$, $^{15}N$]-p53 R249S with and without 70 μM [U-$^2H$]-DNAJA2 (~15% DNAJA2-p53 complex) in 50 mM NaPi pH 7.2, 75 mM KCl, 1 mM DTT, and 0.02% $NaN_3$ buffer. A constant-time relaxation delay of 25 ms was employed with CPMG frequencies ranging between 40 and 1000 Hz.

##### Microsecond relaxation rates measurements

$R_{1\rho}(2H'_zN_z)$, $R_{1\rho}(2H_zN'_z)$, $R_{1\rho}^2(2H'_zN'_z)$, and $R_1(2H_zN_z)$ relaxation rates were measured on a sample containing 360 μM [$^2$H, $^{15}$N]-p53 R249S with and without 70 μM [U-$^2$H]-DNAJA2 (~15% DNAJA2-p53 complex) in 50 mM NaPi pH 7.2, 75 mM KCl, 1 mM DTT, and 0.02% $NaN_3$ buffer. The NMR experiments were measured at 28ºC on a static magnetic field of 14.1T (600 MHz proton frequency) using pulse schemes for measuring the rotating frame relaxation rates [36]. The relaxation rates were measured with delays up to 20 ms, and $^1$H and $^{15}$N continuous wave (CW) spin-lock field strengths were applied with 12.5 kHz and 2 kHz, respectively. $R_2(2H_xN_z)$ rate is derived from the related rotating frame relaxation rate $R_{1\rho}(2H'_zN_z)$ that is measured in the presence of a continuous $^1$H spin-lock radio frequency field, $R_2(2H_zN_x)$ follows from $R_{1\rho}(2H_zN'_z)$. At the same time, $R_2(2H_xN_x)$ is determined from $R_{1\rho}^2(2H'_zN'_z)$ measured in the presence of a double $^1$H-$^{15}$N spin-lock.

All relaxation rates were determined by fitting a single exponential decay function, $I(T_{relax}) = A\exp(-RT_{relax})$, to the measured intensity versus $T_{relax}$ profile. Microsecond chemical exchange contributions, $R_{ex,\mu s}$, were calculated using the following equation -

$$R_{ex,\mu s}(N_x) = \frac{1}{2}\left\{R_1(2H_zN_z)\left(-1+\frac{4c_N^2}{3d_{HN}^2}\right) + R_2(2H_zN_x)\left(1-\frac{4c_N^2}{3d_{HN}^2}\right) + R_2(2H_xN_z)\left(-1-\frac{4c_N^2}{3d_{HN}^2}\right) + R_2(2H_xN_x)\left(1+\frac{4c_N^2}{3d_{HN}^2}\right)\right\}$$

where $d_{HN} = (\mu_0/4\pi)\hbar\gamma_H\gamma_N r_{HN}^{-3}$, $c_N = B_0\gamma_N\Delta\sigma_N\sqrt{((1+\eta_N^2/3)/3)}$, $\mu_0$ is the permeability of free space, $\hbar$ is Planck's constant divided by 2π, $\gamma_H$ and $\gamma_N$ are the gyromagnetic ratios of $^1$H and $^{15}$N, respectively, $r_{HN}$ is the vibrationally averaged distance between $^1$H and $^{15}$N nuclei, $B_0$ is the static magnetic field strength, $\Delta\sigma_N = \sigma_{11,N} - (\sigma_{22,N} + \sigma_{33,N})/2$ (shift anisotropy), and $\eta_N = (\sigma_{22,N} - \sigma_{33,N})/(\sigma_{11,N} - \sigma_{iso,N})$ (reduced asymmetry), where $\sigma_{11,N}$, $\sigma_{22,N}$, and $\sigma_{33,N}$ are the principal components of the nitrogen CSA tensor [92,93] and $\sigma_{iso,N} = 1/3(\sigma_{11,N} + \sigma_{22,N} + \sigma_{33,N})$.

Uncertainties in relaxation rates were obtained from the covariance matrix method [94] and propagated with a Monte Carlo procedure.

##### Hydrogen-deuterium (H/D) exchange

H/D exchange experiments were conducted on 100 μM $^{15}$N-labeled p53 WT or R249S mutant in 50 mM NaPi pH 7.2, 75 mM KCl, 1 mM DTT, and 0.02% $NaN_3$ buffer. Progress of the exchange process between amide protons and deuterium was followed by collecting a series of successive HSQC spectra starting immediately after the sample resolubilization in $D_2O$. All exchange experiments were conducted on a 1000 MHz Bruker Avance NEO spectrometer at either 25°C or 37°C. The first HSQC spectrum was collected after ~20 min, and the rest of the spectra were acquired at a 30 min intervals for 24 hours.

The signal intensities $I(t)$ of each $^1$H-$^{15}$N cross-peak at time $t$ after start of the H/D exchange were fitted to the equation $I(t) = I_0\exp(-k_{ex}t)$, in which $I_0$ is the initial signal intensity and $k_{ex}$ is the H/D exchange rate.

#### Aggregation prevention

All chaperone activity assays were completed using a 96-well plate and a BioTek Synergy H1 or Infinite 200 PRO Tecan plate reader.

##### p53 aggregation

The auto-aggregation of p53 R249S and R282W variants was monitored by following the increase in light scattering in 50 mM NaPi pH 7.2, 150 mM NaCl buffer. The aggregation assay was performed at 37 °C with continuous shaking (567 rpm) and monitored at 400 nm in flat bottom, clear, 96-well plates (Nunc) sealed with optical adhesive film (Applied Biosystems). Samples were run in triplicates and the experiments were repeated at least 3 times with similar results.

##### Insulin aggregation

Human insulin (40 µM; Sigma-Aldrich) was dissolved in 2.5% acetic acid and dialyzed overnight at 4°C to 50mM PBS pH 7.4 buffer. Aggregation was initiated by the addition of 2 mM DTT. The aggregation was monitored at 37 °C by measuring light scattering at 400 nm as a function of time. DNAJA2 variants were added to a final concentration of 40 µM. Assays were acquired in an area scan mode with a 3x3 matrix for each well in clear, flat-bottom, 96-well plates (Nunc) sealed with optical adhesive film (Applied Biosystems).

##### Tau aggregation

Human tau 2N4R (10 µM) was pre-incubated in the presence or absence of DNAJA2 (40 µM) or DNAJA2$^{\Delta BH}$ (40 µM) for 10 minutes at 37 °C. All proteins in the assay were buffer exchanged into the assay buffer (50 mM HEPES pH 7.4, 50 mM KCl, and 2 mM DTT). Thioflavin T (ThT; Thermo Scientific) at a final concentration of 10 µM was added and the aggregation was induced by the addition of a 5 µM freshly prepared heparin salt solution (Sigma). Aggregation reactions were run at 37 °C with continuous shaking (567 rpm) and monitored by ThT fluorescence (excitation = 440 nm, emission= 485 nm), using an area scan mode with a 3x3 matrix for each well. Black, flat-bottom, 96-well plates (Nunc) sealed with optical adhesive film (Applied Biosystems) were used. The experiments were conducted in triplicate and the mean ± standard deviation is reported.

#### Sedimentation assay

To quantify the amount of insoluble / aggregated p53, samples were centrifuged at 17,000*g* for 15 min. The supernatant and pellet fractions were loaded separately on an 12% SDS page and quantified by immunoblotting (p53 pab240 antibody, Abcam). Immunoblots were imaged by ChemiDoc XRS+ system (Bio-Rad).

#### Fluorescence anisotropy measurements

Steady-state equilibrium binding of DNAJA2 variants to p53 DBD WT and R249S and R282W destabilizing mutants was measured by fluorescence anisotropy using 100 nM of fluorescently tagged DNAJA2 (C280- AF488) or DNAJA2$^{mono}$ (C280- AF488). p53 misfolding was induced by heating the samples for 30 min at 37 °C and data was acquired on a BioTek Synergy H1 plate reader in black, flat-bottomed 384-square-well plates. The excitation filter was centered on 485 nm with a bandwidth of 20 nm, and emission filter was centered on 528 nm with a bandwidth of 25 nm. Data were fit to a one-site binding model using GraphPad Prism.

#### Thermal melts

Thermal melts were performed with a nanoDSF instrument (NanoTemper), which was used to monitor the intrinsic fluorescence of p53 DBD WT and R249S or R282W mutants as a function of temperature. Capillaries contained ~10 µL of each protein in a 50 mM NaPi pH 7.2 buffer supplemented with 150 mM NaCl and 2.5 mM DTT. The initial temperature was 20 °C and was set to increase by 1 °C per minute. Fluorescence readings were recorded at 330 and 350 nm, and the melting temperature ($T_m$) was extrapolated from the inflection point.

##### Hsp70 ATPase activity determination

Hsp70 phosphate-release rates after ATP hydrolysis were measured under steady-state conditions by monitoring the change in fluorescence of the phosphate-binding protein (PBP) A197C mutant, which was labelled at Cys197 using 7-diethylamino-3-[*N*-(4-maleimidoethyl)carbamoyl]coumarin (MDCC, CDX-D0198 from Adipogen). Fluorescence was measured in a Synergy H1 plate reader by exciting at 430 nm and measuring at 465 nm.
All reactions contained PBP, 0.25 µM HSP70 in 50 mM HEPES pH 7.5, 25 mM KCl, 10 mM $MgCl_2$ and 2 mM DTT, and 0.25 µM of DNAJA2 or DNAJA2$^{\Delta BH}$ mutant. After the plate was incubated at 37 °C for 10 min, the reactions were started by injection of ATP to a final concentration of 100 µM to each well. Wells were then read every 40 s for the first 20 min, and every 2 min for the next 40 min. For each plate, a series of five phosphate concentrations with PBP alone was measured to generate a calibration curve, which was used to correlate fluorescence to concentration of the released phosphate. All ATPase assays were performed in triplicate.

##### Firefly luciferase refolding

Recombinant firefly luciferase (0.2 µM) was incubated for 40 min at 30 °C in denaturation buffer (25 mM HEPES-KOH, pH 7.5, 50 mM KCl, 10 mM $MgCl_2$, 2 mM DTT, 6 M guanidinium chloride). To start the refolding reaction, the denatured luciferase was diluted 150-fold into refolding buffer (25 mM HEPES, pH 7.5, 50 mM KCl, 10 mM $MgCl_2$, 1 mM DTT, 0.1 mg $ml^{-1}$ BSA, 1 mM ATP, 20 mM creatine phosphate, 6 µg creatine kinase) supplemented with the chaperones and incubated at 30 °C. Each refolding reaction contained 1 µM of indicated DNAJA2 variants, 2 µM Hsc70, and 0.1 µM Hsp110. Luminescence was measured after 120 min by addition of 50 µM of luciferase reagent (Promega) to 9 µl of the refolding reaction on in a white 96 well (Nunc) plate using a Synergy H1 plate reader (BioTek).

#### Cell culture and transient transfection.

###### SaOS-2 Cells

Human osteosarcoma SaOS-2 (p53-null) cell line (ATCC HTB-85) were maintained in DMEM supplemented with 10% FCS (Gibco), 4 mM L-glutamine, 1% penicillin-streptomycin in a 5% $CO_2$ humidified incubator at 37 °C.
Cells were seeded at $1x10^5$ cells per 6-well plate and allowed to reach 90% confluency before transfection. Co-transfections of 1.5 µg pcDNA5/FRT/TO-V5-JDPs (DNAJA2, DNAJB1, DNAJA2$^{\Delta BH}$) and 0.5 µg of pcDNA3.1-p53 were performed using 3.5 µl of Lipofectamine 3000 and 4 µl of P3000 (Invitrogen), according to manufacturer's instructions. The DNA complex, P3000, and Lipofectamine were prepared in 250 µl of OptiMEM and incubated for 20 minutes before addition to plated cells. Cells were analyzed 24h after transfection.

###### PaTu 8988t Cells

TP53 homozygous mutant (p.Arg282Trp) human pancreatic cancer cell line PaTu 8988t were grown in DMEM supplemented with (v/v) 10% FCS and Pen-Strep (BI, 03-031-1B).
For transfection, cells were seeded at $7x10^4$ cells per 24-well plate and were allowed to reach 90% confluency before transfection. Depletion of DNAJA1 by siRNA (40 nM, Dharmacon ON-TARGETplus Human DNAJA1 siRNA SMARTpool) and overexpression of DNAJA2 WT or variants (0.5 µg) were performed simultaneously using Lipofectamine 3000 (Invitrogen). Scrambled RNA (Non-targeting Pool, Dharmacon) and an empty pcDNA5.1 vectors were used as controls. The transfection media was replaced 6 h post transfection and the cells were analyzed 48-72 h later.

**Western blots**

SaOS-2 cells were lysed in TBS with 1% CHAPS, benzonase and protease inhibitor cocktail (Roche). For SDS-PAGE, whole cell lysates were denatured at 95 °C for 10 min in Laemmli sample buffer and then loaded on ExpressPlus PAGE gel (GenScript). For Blue Native-PAGE, cell lysates were supplemented with 20% glycerol and 5 mM Coomassie G-250 before loading onto 3–12% Novex Bis-Tris gradient gels. Electrophoresis was performed in a running buffer containing 50 mM BisTris and 50 mM Tricine (plus 0.004% Coomassie G-250 in cathode buffer) under fixed voltage (100 V) for 2 hours. Proteins were transferred from gels onto PVDF membranes and fixed with 8% acetic acid for 30 minutes. The membranes were subsequently blocked with 3% BSA in TBS at 4 °C for 1 h and incubated with primary antibodies against the indicated proteins followed by appropriate secondary antibodies conjugated with fluorophores. All blots were analyzed with an ImageQuant LAS 4000 biomolecular Imager (GE Healthcare) and quantified in ImageJ (NIH).

The following antibodies were used: p53 (DO-1, sc-126, Santa Cruz Biotechnology, 1:500 dilution), V5-DNAJA2 (V5 probe-E10, sc-81594, Santa Cruz Biotechnology, 1:500 dilution), DNAJA2 (PA5-65583, Invitrogen, 1:1000 dilution), DNAJA1 (sc-59554, Santa Cruz Biotechnology, 1:500 dilution), and Actin (MA5-11869, Invitrogen, 1:500 dilution).
PaTu 8988t cells were lysed on ice using lysis buffer (50 mM Tris–HCl, pH 7.4, 150 mM NaCl, 1% NP-40, 1 mM EDTA) supplemented with protease inhibitor cocktail (Roche). Protein concentrations were determined using a BCA Protein Assay Kit (Thermo Fisher Scientific). Aliquots of lysates were solubilized in Laemmli sample buffer, and equal amounts of proteins were separated on 4-20% gradient SDS-PAGE gels (GenScript). Proteins were transferred onto nitrocellulose membranes and then blocked with 5% nonfat dry milk in TBS buffer for 1 hour. The membranes were then incubated with primary antibodies, specific to the protein of interest, in TBS overnight at 4 °C. After incubation with the appropriate secondary antibody conjugated with HRP, ECL (Clarity, Bio-Rad) was used for protein detection. Immunoblots were obtained using the ChemiDoc MP Imaging System (Bio-Rad). Densitometry was measured with ImageJ software (NIH).

The following antibodies were used: p53 (DO-1, sc-126, Santa Cruz Biotechnology, 1:1000), DNAJA1 (Abcam, ab192904, 1:1000), V5 (Abcam, ab206566, 1:1000), DNAJA2 (Abcam, ab157216, 1:1000), vinculin (Abcam, ab129002, 1:1000), Actin (MA5-11869, Invitrogen, 1:500), goat anti mouse IgG-HRP (JIR 115-035-003), and goat anti rabbit IgG-HRP (JIR 111-035-003).

**Supernatant/pellet fractionation**

To quantify the amount of soluble/insoluble mutant p53 in SaOS2 cells upon co-overexpression of JDPs, the cell lysate was centrifuged at 12,500g for 15 minutes at 4 °C and fractionated into supernatant and pellet. The fractions were subsequently analyzed by SDS-PAGE and western blot detection of p53 (DO-1, sc-126, Santa Cruz Biotechnology, 1:500 dilution) or actin (MA5-11869, Invitrogen, 1:500 dilution) as control. Relative abundance of p53 in the pellet was calculated from densitometry analysis with Image J (NIH).

**Immunofluorescence staining**

SaOS-2 cells were plated 24 hours after transfection on standard coverslips. The coverslips were washed twice with phosphate-buffered saline (PBS) and fixed in 4% (v/v) formaldehyde for 15 minutes at RT. Cells were then permeabilized and blocked with 0.1% (w/v) Triton X-100 and 3%(w/v) BSA in PBS for 30 minutes. The primary antibodies for p53 (DO-1 mouse monoclonal, Sata Cruz Biotechnology, dilution 1:500) and V5-tagged JDPs (V5 rabbit polyclonal, Cell Signalling, dilution 1:1000) were incubated overnight at 4 °C. Secondary antibodies (goat anti-mouse Alexa 568 or goat anti-rabbit Alexa488, Invitrogen, dilution1:500) were incubated for 1 h at RT followed by staining with DAPI (Invitrogen, dilution 1:5000). Finally, coverslips were treated with antifade reagent (ProLong Gold, Invitrogen) and kept in the dark for 24 h before image acquisition with a SP5 confocal fluorescence microscope (Leica) or SP8 STED 3X microscope with FALCON FLIM (Leica).

PaTu 8988t cells were plated on a standard poly-D-lysine-coated glass coverslip. The coverslips were fixed in 3.7% (v/v) formaldehyde in PBS for 15 minutes, permeabilized for 2 minutes with 0.1% Triton in PBS, and blocked with Blocking Buffer (0.1% Tween and 5% (w/v) BSA in PBS) for 1 h. Cells were then incubated with primary anti-V5 tag antibody (Abcam, 1:200) and secondary antibody (goat anti-rabbit Alexa Fluor 488, Abcam, 1:2000). Coverslips were then mounted with antifade reagent (Fluoroshield™ with DAPI, Sigma-Aldrich) and kept in the dark for 24 h. Images were acquired on an Olympus IX51 microscope equipped with Olympus XM10 camera, and using the 100X magnification objective and analysed using ImageJ software (NIH).

**Foci quantification**

For the quantification of the fraction of cells containing aggregated p53, slides were imaged on a SP5 confocal fluorescence microscope (Leica) at 40x magnification, 24 hours after transfection. To allow the detection of cytoplasmic foci across different experimental conditions, the intensity was adjusted to avoid saturation of fluorescence signals, leading to minor differences in microscopy settings. Representative images were acquired using the 60x magnification objective for clarity. The images were subsequently processed using ImageJ software (NIH).
Transfected SaOS-2 cells, positive for p53 and JDP immunofluorescence staining were scored for the presence of p53 aggregates (bright fluorescent foci) in the cytoplasm. For all experimental conditions, 3 biological replicates were performed with ~100 cells scored per replicate in Fig.1 and ~300 cells per replicate in Fig. 3.

**Co-immunoprecipitation**

Cells were lysed in TBS with 1% CHAPS, benzonase, and protease inhibitor cocktail (Roche) for 30 min on ice. The cell lysate was subsequently incubated with anti-V5 antibody (V5 rabbit polyclonal, Cell Signalling, dilution 1:1000) overnight at 4°C. Then 15 µL of protein G Dynabeads (Thermo Fisher Scientific) was added. After incubation at RT for 2 hours, the beads were rinsed five times with 500 µL TBS, and subsequently eluted by heating at 95 °C in presence of SDS. Input and IP samples were analyzed by SDS-PAGE and western blot.

**Cycloheximide Chase**

The translation inhibitors cycloheximide (CHX, Sigma–Aldrich) was used to evaluate the kinetics of endogenous p53 mutant degradation. 72 h post transfection, PaTu 8988t cells were treated with 100 µg/ml (final concentration) cycloheximide. Immediately after treatment and after 4, 10, 13, 16, and 20 hours of incubation with CHX, the cells were lysed, and the cellular lysates were separated on a 4-20% gradient SDS-PAGE gels (Genscript) and blotted against p53 (DO-1, sc-126, Santa Cruz Biotechnology, 1:1000). Total protein levels were quantified and adjusted by WB using vinculin (Abcam, ab129002, 1:1000) as loading control.

In order to validate mutant p53 (p.Arg282Trp) proteasomal degradation, DNAJA1-depleted PaTu 8988t cells were treated with 30 µM MG-132 (Cell Signaling), simultaneously with CHX treatment (4-20 h). DMSO was used as control.

The amount of protein at the start of the chase (time 0) was set to 100%, and the amounts of protein at subsequent time points were expressed as a percent of the starting material. The experiments were repeated three times.

##### Quantitative Reverse Transcription-PCR (RT-qPCR)

The effects of p53 mutants on transactivation of target genes (MDM2 and p21) was determined by RT-qPCR 24 hours after transfection. Total RNA was isolated from cells using RNeasy mini kit (QIAGEN). Reverse transcription was performed using the iScript cDNA Synthesis Kit (Bio-Rad) according to the manufacturer's protocols. Quantitative real-time PCR was performed using TB Green Premix Ex Taq II (Takara) on LightCycler 480 II real-time PCR detection system (Roche). ACTB was used as a reference gene.

The effect of DNAJA2 variants on the oncogenic PLK2 gene mRNA levels was quantified in PaTu 8988 cells. Total RNA was isolated from cells using RNeasy mini kit (QIAGEN), 60 hours post-transfection. Reverse transcription was performed using the qScript cDNA Synthesis Kit (QuantaBio) according to the manufacturer's protocols. Quantitative real-time PCR was performed using Fast SYBR Green Master Mix (Applied Biosystems) on StepOnePlus Real-Time System (Applied Biosystems). GAPDH was used as a reference gene.

A list of the primer sequences used for qPCR in this study is provided in Supplementary Table 1.

#### Characterization of p53-DNAJA2 complexes

###### Negative stain electron microscopy

p53 R249S or R282W aggregates were generated by incubating 10 µM of mutant p53 DBD for 3 h at 37 °C with shaking (800 rpm) in 50 mM NaPi pH 7.2, 150 mM NaCl, 2.5 mM DTT. p53-DNAJA2 complexes were prepared similarly, but 40 µM DNAJA2 was added to p53 prior to incubation at 37 °C and the complexes were isolated on a SEC column (200 Increase 10/300 GL, GEHealthcare). DNAJA2-p53 complexes or p53 aggregates (3.5 µl) were deposited on glow discharged carbon-coated copper EM grids (Electron Microscopy Sciences), stained with three drops of 2% w/v Uranyl-acetate, and air-dried. Imaging was performed on an FEI Tecnai T12/T12 Spirit transmission electron microscope at 120kV and a magnification of 1650-30000 times.

###### SEC-MALS

p53-DNAJA2 complexes were prepared by incubating 30 µM p53 and 120 µM DNAJA2 at 37 °C for 3 hours. The complexes were then isolated on a SEC column (200 Increase 10/300 GL, GEHealthcare), and 100 µL were reinjected into a pre-equilibrated Superdex 200 Increase 10/300 GL column coupled to a multi-angle light scattering detector (DAWN HELEOS II, Wyatt) and refractometer (Optilab T-rEX, Wyatt). Data were analyzed using the ASTRA software (Wyatt Technology).

###### Dynamic Light Scattering (DLS)

Samples containing 10 µM R249S and 20, 30, 40, 60, or 100 µM DNAJA2 were incubated for 3 hours at 37°C with shaking (800 rpm) in 50 mM NaPi pH 7.2 buffer supplemented with 150 mM NaCl and 2.5 mM DTT. After the incubation period, the complexes were isolated by size exclusion chromatography (Superdex 200 Increase 10/300 GL, GE Healthcare). The fractions consisting of the p53-DNAJA2 complex were subjected to dynamic light scattering (DLS) measurements performed by DynaPro Plate Reader III (Wyatt) using a laser at wavelength 830 nm and operating with a 158° scattering angle. Thermo Fischer 96 well plates (265301, Nunc) were used, with a sample volume of 200 µL per well. For each sample, 5-10 reads were acquired and processed and the data were analyzed using the accompanying software Dynamics, version 7.10.1.21.

**Release of monomeric p53 by Hsp70 system**

Samples containing 2 µM p53 WT, R249S, or R282W DBDs were incubated for 3 hours at 37 °C with shaking (800 rpm) either alone or with 20 µM DNAJA2 variants. A total volume of 0.5 ml was incubated in 50 mM HEPES pH-7.1, 150 mM KCl, 2.5 mM DTT buffer. The different components of the Hsp70 system (40 µM Hsp70, 20 µM DNAJB1, and 2 µM Hsp110) were then added to the reaction mixtures to a final volume of 1 ml. The reaction was allowed to continue for 16 h at 25 °C with a continuous supply of ATP through dialysis in 50 mM HEPES pH-7.1 buffer supplemented with 150 mM KCl, 10 mM $MgCl_2$, and 2.5 mM DTT. Next, the samples were centrifuged for 10 min at 17000 RCF to separate the insoluble p53 aggregates, and the supernatant was subjected to SEC fractionation (Superdex 200 Increase 10/300, GE Healthcare) in 50 mM HEPES pH-7.1, 150 mM KCl, 2.5 mM DTT buffer.

The SEC fractions corresponding to elution volumes of 8-20 ml, and the insoluble fractions separated by centrifugation prior, were run on a 12% SDS-PAGE and blotted against p53 (pab240, Abcam). Monomeric folded p53 protein (2 µM p53 DBD) which eluted in fractions 18 and 19 was used as a control. Gel filtration standards (#1511901, Bio-Rad) were used for the determination of molecular weight distribution.

**Electrophoretic mobility shift assay (EMSA)**

p53 DBD WT, R249S, or R282W variants were incubated for 20 minutes with 500 nM of Cy5.5-labeled DNA-oligo p21 (5′TGGCCATCAGGAACATGACCCAACATGTTGAGCTCTGGCA′3) in 50 mM HEPES pH-7.1, 150 mM KCl, 2 mM DTT buffer, at 25 °C. p53-DNAJA2 complexes, p53 aggregates, or chaperone-treated p53 samples were generated as described previously. Samples were run on a 7% native polyacrylamide gel in Tris-Glycine buffer with 0.1 mM EDTA at 135 V at 4°C in the dark. The gels were imaged using a Li-Cor Scanner (Odyssey Li-Cor Biosciences) at 700 nm and analysed using Image Studio v5.2 software.

#### Supplementary Figure Legends

**Figure S1. Class A JDPs suppress mutant p53 misfolding and aggregation in vitro and in cell lines. (A)** Sedimentation assays for R282W and R249S p53 mutants incubated for 5 hours at 37°C in the absence or presence of DNAJA2. The soluble (S) and insoluble (P) fractions were blotted against p53. **(B-D)** $^{1}H$-$^{15}N$ HSQC NMR spectra of $^{15}N$ labeled p53 R249S DBD alone (black), and with 2-fold molar excess of unlabeled DNAJB1 (**B**), DNAJA1 (**C**) or DNAJA2 (**D**) recorded at 37°C. No changes to the spectrum are detected upon addition of DNAJB1, while addition of both DNAJA1 and DNAJA2 caused significant peak broadening, indicating binding to class A, but not to class B JDPs. **(E)** Melting curves (first derivatives) showing the stability of WT (blue), R249S (pink), and R282W (orange) p53 proteins. **(F)** Representative images of SaOS2 p53 null cells co-overexpressing WT p53 and V5-DNAJA2 or V5-DNAJB1 immunostained for p53 (red) and V5 (green) and quantified in Figure 1G. Crop images are the overlay of the p53 signal with DAPI. Scale bar: 10 µm. WT p53 shows a predominantly nuclear localization that is unaffected by JDP overexpression. **(G)** Blue-Native PAGE of p53 WT, R249S, and R282W mutant stained with Do-1 p53 antibody. While WT p53 is found primarily in a tetrameric form, the destabilized p53 mutants are present as higher MW species, reflecting aggregation. **(H)** Western blots showing the protein levels of WT p53, R249S and R282W co-expressed with V5-DNAJA2 or V5-DNAJB1 shown in Figures 1F,G. **(I-J)** Representative results of the supernatant/pellet fractionation of cell lysate expressing R249S **(I)** or R282W **(J)** p53 co-expressed with the empty vector, DNAJA2, or DNAJB1. DNAJA2 overexpression results in a significant reduction of p53 in the pellet fraction (59% and 45% respectively, n=3, $p<0.05$) for both mutants.

**Figure S2. DNAJA2 binds to structured β-strands at the core of p53 DBD. (A-B)** Residue-resolved NMR signal attenuation ($I/I_0$) of R249S (A) or R282W (B) p53 mutants upon addition of 2-fold excess of deuterated DNAJA2. Decrease in intensity indicates binding, which is mainly localized to β-strands 4, 8 and 9. **(C)** Fluorescence anisotropy binding assays of Alexa Fluor-488 labeled DNAJA2, titrated with increasing concentrations of p53 WT (blue), R249S (pink), or R282W (orange), measured at 28°C. R249S and R282W p53 mutants bind DNAJA2 with 55.9±2.6 µM and 6.3±0.7 µM affinity, while no significant binding was detected for WT p53. Data are means ± SEM (n = 2). **(D)** CPMG RD data showing the difference between the effective $R_2$ rates at high (1 kHz) and low (40 Hz) CPMG frequencies for 15% DNAJA2-p53 complex. The fact that no significant $\Delta R_{2,eff}$ is observed suggests the absence of chemical exchange contribution in the ms timescale. **(E-F)** Residue-specific microsecond chemical exchange contributions, $R_{ex,\mu s}$, derived from the four relaxation rates $R_2(2H_xN_z)$, $R_2(2H_zN_x)$, $R_2(2H_xN_x)$, and $R_1(2H_zN_z)$ for free p53 R249S (E) and p53 R249S in complex with 15% DNAJA2 (F). The error bars include both experimental errors and errors introduced by the uncertainty in $\Delta\sigma N$ [37]. A clear increase in microsecond chemical exchange is visible in p53 β4 and β9 regions only upon DNAJA2 addition.

**Figure S3. DNAJA2 recognizes the increased dynamics in the destabilized DBD of mutant p53s.**
**(A)** Hydrogen/Deuterium (H/D) exchange NMR curves for residues located in β4 (V157 & A159), β8 (I232 & Y234), and β9 (T253 & I255) in p53 WT (blue) and R249S (pink) DBDs. **(B)** Residue-specific intensity measured 2 hours into the H/D exchange for p53 WT (blue) and R249S (pink). **(C)** Residue-specific intensity 2 hours into the H/D exchange for p53 WT measured at 28°C (dark blue) or 39°C (light blue).

**Figure S4. DNAJA2 β-hairpin region is required for p53 aggregation prevention, but not for other DNAJA2 chaperoning functions. (A-B)** Intensity changes ($I/I_0$) of $^2H$, $^{15}N$ labeled monomeric DNAJA2 homolog from *S.cerevisiae* (Ydj1$^{111\text{-}351}$) upon binding to R249S (A) or R282W (B). p53 binds to a β-hairpin insertion in the zinc finger-like region (ZFLR). **(C-D)** Chemical shift perturbation plots of $^2H$, $^{15}N$ labeled monomeric DNAJA2 homolog from *S.cerevisiae* (Ydj1$^{111\text{-}351}$) upon binding to p53 core β4 (C) and β9 peptides (D). The unfolded peptides interact with the client-binding CTDI and CTDII domain in DANJA2, and not to the newly identified β-hairpin site. **(E)** $^1H$ - $^{15}N$ HSQC NMR spectra of $^{15}N$ labeled p53 R249S DBD alone (black), and with 2-fold molar excess of DNAJA2 (red) or DNAJA2$^{\Delta BH}$ (violet). Significant peak broadening is detected upon addition of wild-type DNAJA2, while no binding is detected with the DNAJA2$^{\Delta BH}$ mutant. **(F)** Sedimentation assays for R282W and R249S p53 mutants incubated for 5 hours at 37°C in the absence and presence of DNAJA2 WT or DNAJA2$^{\Delta BH}$ mutant. The soluble (S) and insoluble (P) fractions were blotted against p53. **(G)** ThT-based aggregation assay of 10 μM tau in the presence of 40 μM DNAJA2 (red) or DNAJA2$^{\Delta BH}$ (violet). Similar aggregation-prevention activity was observed for the wild-type and mutant JDPs. Data are means ± SEM (n = 3). **(H)** Enhancement of Hsp70 steady-state ATPase activity by wild-type DNAJA2 (red) or DNAJA2$^{\Delta BH}$ mutant (violet) JDPs. The basal activity of Hsp70 is shown in grey. Both DNAJA2 constructs activate the Hsp70 chaperone to the same extent. Data are means ± SEM (n = 3). **(I)** Refolding of heat-denatured firefly luciferase by a chaperone refolding system consisting of Hsp70, NEF, and WT DNAJA2 (red) or mutant DNAJA2$^{\Delta BH}$ (violet). DNAJA2 lacking the β-hairpin shows high protein refolding yields. Data are means ± SEM (n = 5).

**Figure S5. DNAJA2 β-hairpin site is essential for p53 aggregation-prevention in cells. (A)** Colocalization of p53 R249S co-expressed with V5-DNAJA2 or V5-DNAJA2$^{\Delta BH}$ by stimulated emission depletion (STED) microscopy in SaOS2 cells. Scale bar: 10 μm and 1 μm as indicated. WT DNAJA2 co-localizes with residual p53 aggregates, while no co-localization is observed for DNAJA2$^{\Delta BH}$. The β-hairpin region thus drives DNAJA2 interaction with destabilized p53 in cells. **(B-C)** Immunostaining **(B)** and western blot of proteins levels **(C)** of mutant p53 (R249S, R282W) co-expressed with V5-DNAJA2 or V5-DNAJA2$^{\Delta BH}$, quantified in Figure 3F. The crop images (corresponding to white boxed areas in the merge image) are the overlay of the p53 signal with DAPI. Scale bar: 10 μm. The deletion of the β-hairpin region abolishes the prevention of aggregation activity of DNAJA2. **(D)** Supernatant/pellet fractionation of cell lysate of p53 R249S co-expressed with an empty vector, DNAJA2 and DNAJA2$^{\Delta BH}$. In contrast to WT DNAJA2, co-overexpression DNAJA2$^{\Delta BH}$ does not reduce the fraction of insoluble p53 mutant. Representative of n=3.

**Figure S6. DNAJA2 chaperones assemble into large oligomeric particles upon binding to destabilized p53. (A)** Size exclusion column (SEC) chromatograms of p53 (black) and DNAJA2 at time 0 (dark red) and following 5 hour incubation at 37°C (red dashed line). **(B)** SEC chromatograms following p53-DNAJA2 complex formation at time 0 (black), and following 5h incubation at 37°C (pink). **(C)** SEC-MALS analysis of isolated DNAJA2-p53 complexes separated on a Superdex 200 Increase 10/300 column. The DNAJA2-p53 complexes remain stable, with no visible dissociation of p53 monomers or DNAJA2 dimers. **(D)** Histograms of DLS-measured hydrodynamic radii of isolated R249S-DNAJA2 (1:2 and 1:3) complexes. **(E)** Hydrodynamic radius of DNAJA2-p53 complexes plotted as a function of DNAJA2 dimer concentration, as determined from DLS measurements. p53 concentration was kept constant at 3 μM. **(F)** DLS measurements of the hydrodynamic radius of isolated DNAJA2-p53 complex, p53 R249S DBD, or DNAJA2 measured for 2 hours at 32 °C. The hydrodynamic radius as a function of time for p53 R249S DBD under non-aggregation inducing conditions (25 °C) are shown as a control. The DNAJA2-p53 complex remains stable over time, while p53 R249S DBD without the DNAJA2 chaperone aggregates within 20 minutes. **(G-J)** Representative negative stain electron micrographs of p53 alone or p53-DNAJA2 mixtures following 3-hour incubation at 37°C. Both R249S **(G)** and R282W **(I)** p53 mutants form large amorphous aggregates in the absence of DNAJA2 and monodisperse ~40 nm particles when in complex with the chaperone **(H, J)**. Scale bar of 500 nm is shown on the bottom of each image.

**Figure S7. Class A JDP chaperones protect the oncogenic p53 mutants from degradation. (A-B)** Quantification of mRNA levels of p53 target genes, p21 **(A)** and MDM2 **(B)**, in SaOS2 cells co-expressing p53 variants (WT, R249S, R282W) and V5-DNAJA2 or V5-DNAJA2$^{\Delta BH}$ by qPCR. Data represent mean values ± s.d (n=3). **(C)** Immunostaining of p53 and DNAJA2 in PaTu-8988 cells transfected with DNAJA1 siRNA and expressing DNAJA2$^{WT}$, DNAJA2$^{\Delta BH}$, or DNAJA2$^{QPN}$ variants. Scrambled siRNA was used as control. **(D)** Immunoblot monitoring p53, DNAJA1, DNAJA2, and vinculin levels in PaTu-8988 cells expressing empty vector, DNAJA2$^{WT}$, DNAJA2$^{\Delta BH}$ or DNAJA2$^{QPN}$ mutants with or without knockdown of DNAJA1. **(E)** MG-132 proteasome inhibitor blocks the turnover of endogenous p53 R282W protein, induced by the depletion of class A JDPs (siDNAJA1). Time points following treatment with cycloheximide (CHX) are shown on top. Vinculin and c-jun were monitored as controls.

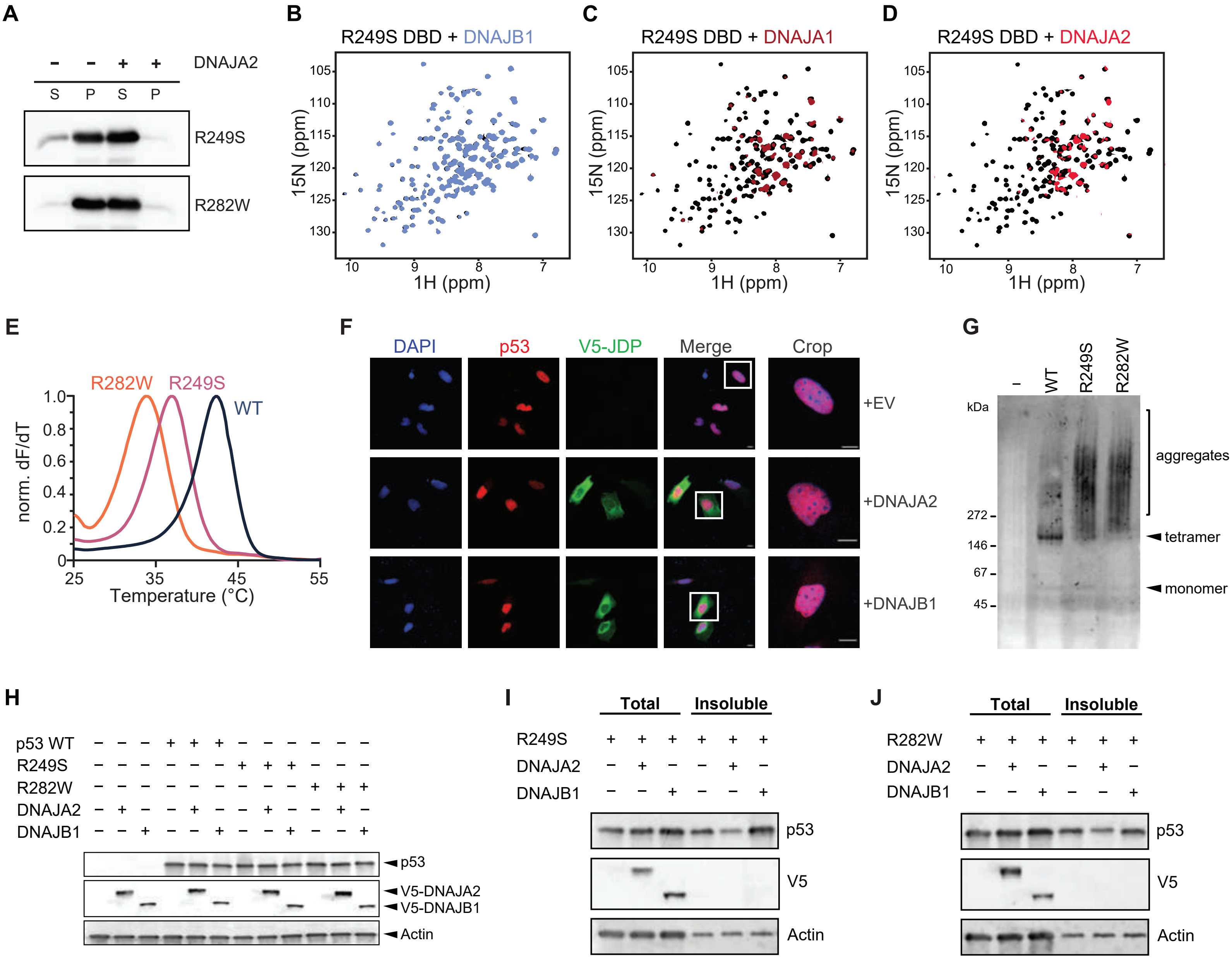

**Figure S1. Class A JDPs suppress mutant p53 misfolding and aggregation in vitro and in cell lines. (A)** Sedimentation assays for R282W and R249S p53 mutants incubated for 5 hours at 37°C in the absence or presence of DNAJA2. The soluble (S) and insoluble (P) fractions were blotted against p53. **(B-D)** $^{1}$H-$^{15}$N HSQC NMR spectra of $^{15}$N labeled p53 R249S DBD alone (black), and with 2-fold molar excess of unlabeled DNAJB1 (**B**), DNAJA1 (**C**) or DNAJA2 (**D**) recorded at 37°C. No changes to the spectrum are detected upon addition of DNAJB1, while addition of both DNAJA1 and DNAJA2 caused significant peak broadening, indicating binding to class A, but not to class B JDPs. **(E)** Melting curves (first derivatives) showing the stability of WT (blue), R249S (pink), and R282W (orange) p53 proteins. **(F)** Representative images of SaOS2 p53 null cells co-overexpressing WT p53 and V5-DNAJA2 or V5-DNAJB1 immunostained for p53 (red) and V5 (green) and quantified in Figure 1G. Crop images are the overlay of the p53 signal with DAPI. Scale bar: 10 µm. WT p53 shows a predominantly nuclear localization that is unaffected by JDP overexpression. **(G)** Blue-Native PAGE of p53 WT, R249S, and R282W mutant stained with Do-1 p53 antibody. While WT p53 is found primarily in a tetrameric form, the destabilized p53 mutants are present as higher MW species, reflecting aggregation. **(H)** Western blots showing the protein levels of WT p53, R249S and R282W co-expressed with V5-DNAJA2 or V5-DNAJB1 shown in Figures 1F,G. **(I-J)** Representative results of the supernatant/pellet fractionation of cell lysate expressing R249S **(I)** or R282W **(J)** p53 co-expressed with the empty vector, DNAJA2, or DNAJB1. DNAJA2 overexpression results in a significant reduction of p53 in the pellet fraction (59% and 45% respectively, n=3, $p<0.05$) for both mutants.

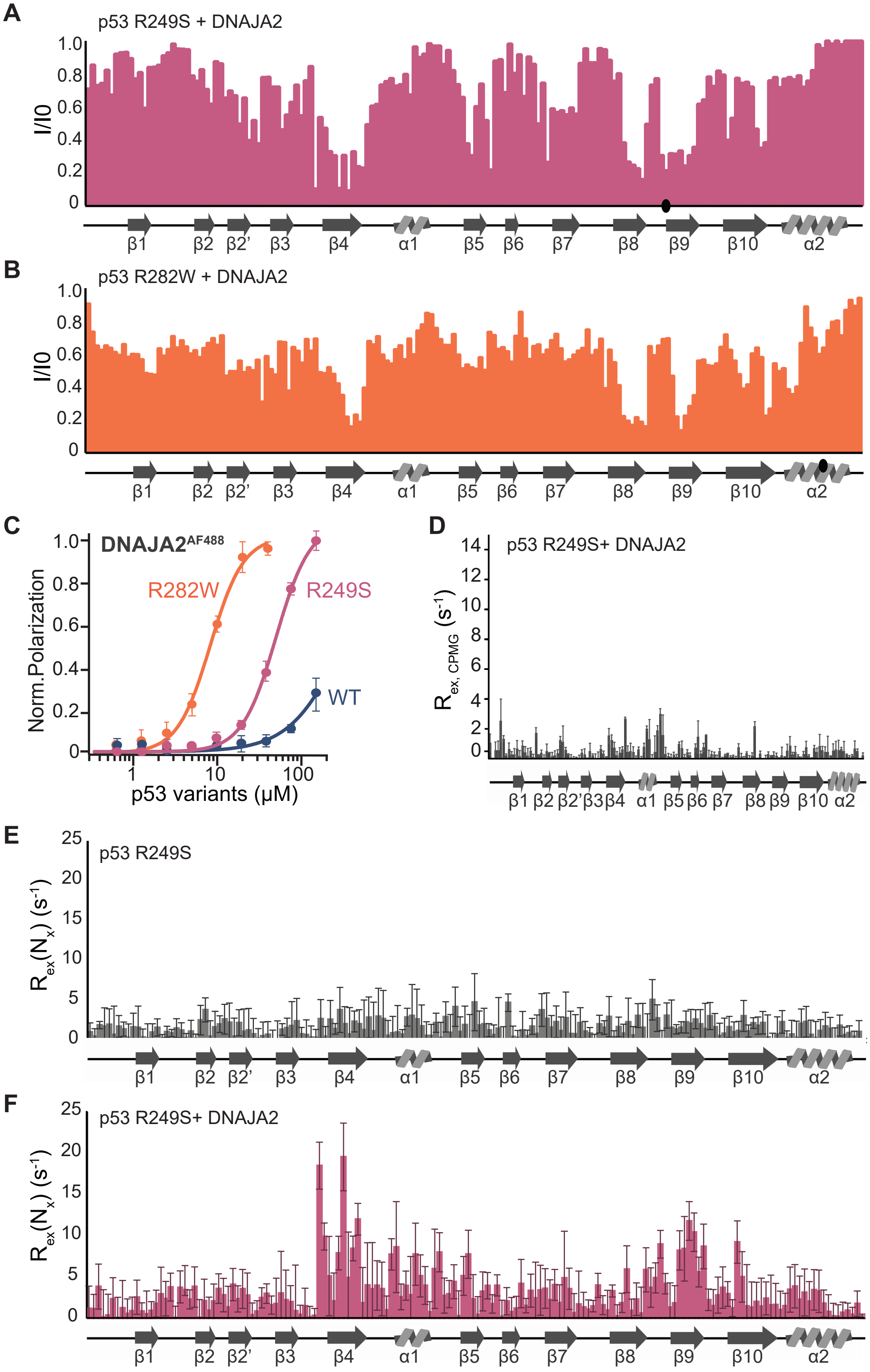

**Figure S2. DNAJA2 binds to structured β-strands at the core of p53 DBD. (A-B)** Residue-resolved NMR signal attenuation ($I/I_0$) of R249S (A) or R282W (B) p53 mutants upon addition of 2-fold excess of deuterated DNAJA2. Decrease in intensity indicates binding, which is mainly localized to β-strands 4, 8 and 9. **(C)** Fluorescence anisotropy binding assays of Alexa Fluor-488 labeled DNAJA2, titrated with increasing concentrations of p53 WT (blue), R249S (pink), or R282W (orange), measured at 28°C. R249S and R282W p53 mutants bind DNAJA2 with 55.9±2.6 μM and 6.3±0.7 μM affinity, while no significant binding was detected for WT p53. Data are means ± SEM (n = 3). **(D)** CPMG RD data showing the difference between the effective $R_2$ rates at high (1 kHz) and low (30 Hz) CPMG frequencies for 15% DNAJA2-p53 complex. **(E-F)** Residue-specific microsecond chemical exchange contributions, $R_{ex,\mu s}$, derived from the four relaxation rates $R_2(2H_xN_z)$, $R_2(2H_zN_x)$, $R_2(2H_xN_x)$, and $R_1(2H_zN_z)$ for free p53 R249S (E) and p53 R249S in complex with 15% DNAJA2 (F). The error bars include both experimental errors and errors introduced by the uncertainty in ΔσN [31]. Clear increase in microsecond chemical exchange is visible in p53 β4 and β9 regions only upon DNAJA2 addition.

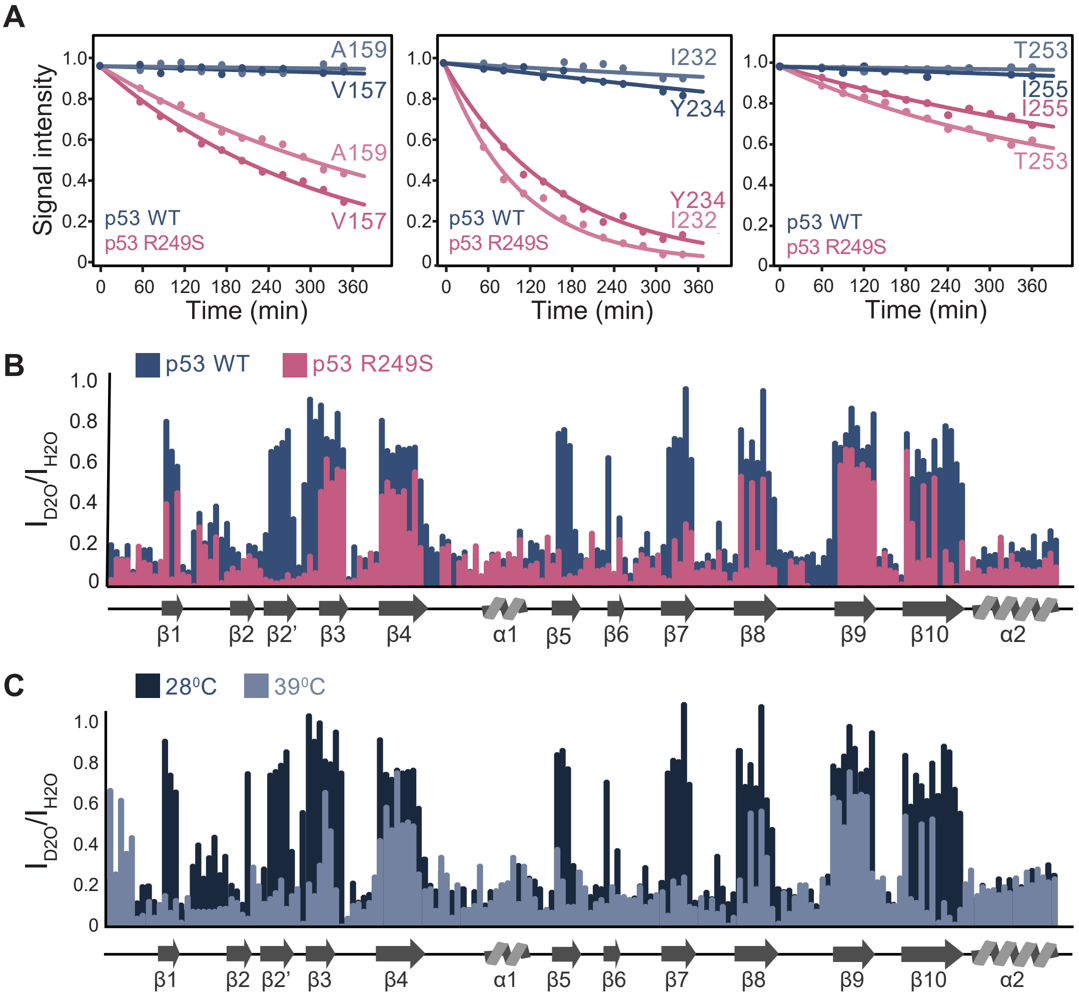

**Figure S3. DNAJA2 recognizes the increased dynamics in the destabilized DBD of mutant p53s (A)** Hydrogen/Deuterium (H/D) exchange NMR curves for residues located in β4 (V157 & A159), β8 (I232 & Y234), and β9 (T253 & I255) in p53 WT (blue) and R249S (pink) DBDs. **(B)** Residue-specific intensity measured 2 hours into the H/D exchange for p53 WT (blue) and R249S (pink). **(C)** Residue-specific intensity 2 hours into the H/D exchange for p53 WT measured at 28°C (dark blue) or 39°C (light blue).

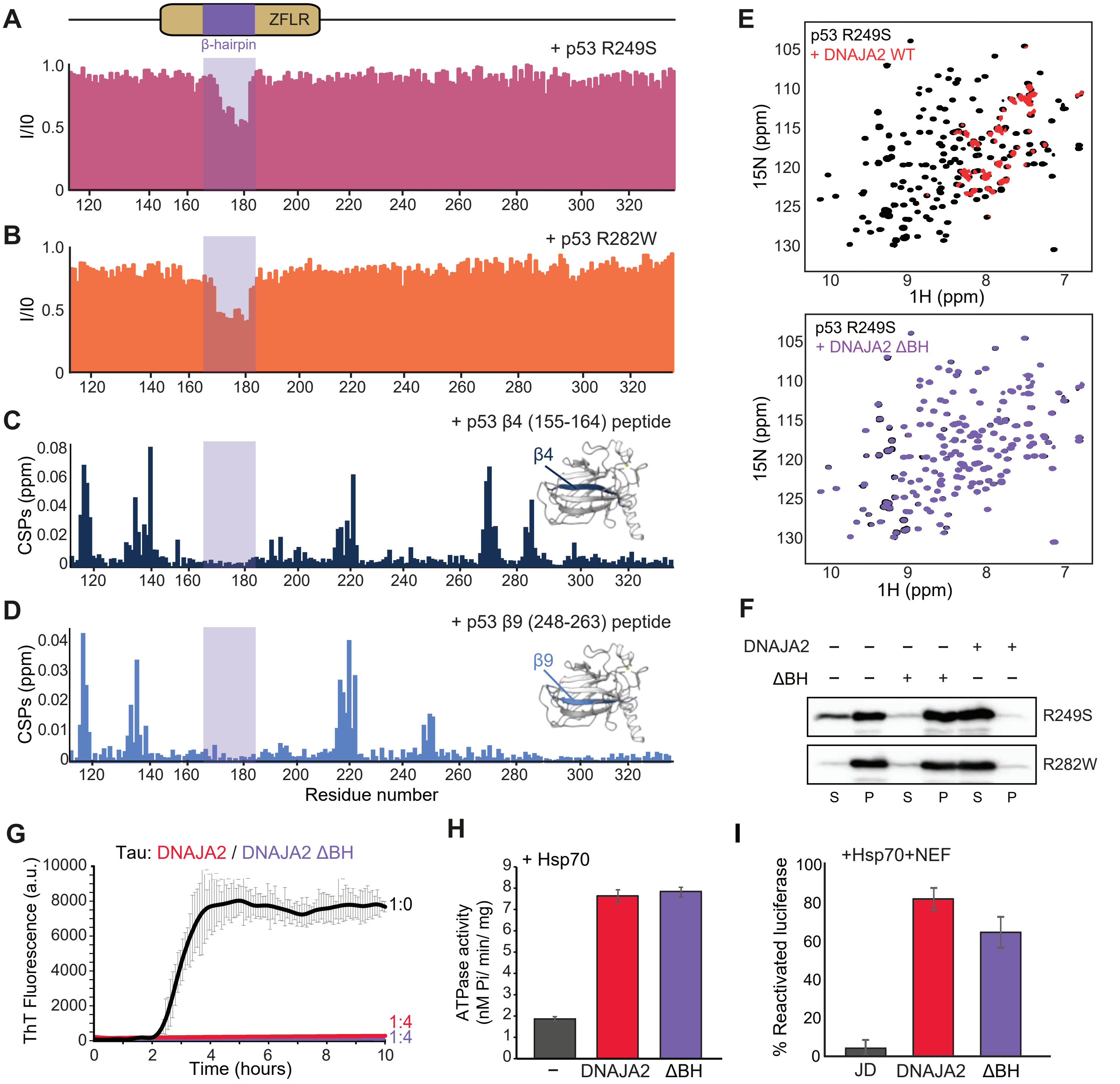

**Figure S4. DNAJA2 β-hairpin region is required for p53 aggregation prevention, but not for other DNAJA2 chaperoning functions. (A-B)** Intensity changes ($I/I_0$) of $^{2}$H, $^{15}$N labeled monomeric DNAJA2 homolog from *S.cerevisiae* (Ydj1$^{111\text{-}351}$) upon binding to R249S (A) or R282W (B). p53 binds to a β-hairpin insertion in the zinc finger-like region (ZFLR). **(C-D)** Chemical shift perturbation plots of $^{2}$H, $^{15}$N labeled monomeric DNAJA2 homolog from *S.cerevisiae* (Ydj1$^{111\text{-}351}$) upon binding to p53 core β4 (C) and β9 peptides (D). The unfolded peptides interact with the client-binding CTDI and CTDII domain in DANJA2, and not to the newly identified β-hairpin site. **(E)** $^{1}$H - $^{15}$N HSQC NMR spectra of $^{15}$N labeled p53 R249S DBD alone (black), and with 2-fold molar excess of DNAJA2 (red) or DNAJA2$^{\Delta BH}$ (violet). Significant peak broadening is detected upon addition of wild-type DNAJA2, while no binding is detected with the DNAJA2$^{\Delta BH}$ mutant. **(F)** Sedimentation assays for R282W and R249S p53 mutants incubated for 5 hours at 37°C in the absence and presence of DNAJA2 WT or DNAJA2$^{\Delta BH}$ mutant. The soluble (S) and insoluble (P) fractions were blotted against p53. **(G)** ThT-based aggregation assay of 10 μM tau in the presence of 40 μM DNAJA2 (red) or DNAJA2$^{\Delta BH}$ (violet). Similar aggregation-prevention activity was observed for the wild-type and mutant JDPs. Data are means ± SEM (n = 3). **(H)** Enhancement of Hsp70 steady-state ATPase activity by wild-type DNAJA2 (red) or DNAJA2$^{\Delta BH}$ mutant (violet) JDPs. The basal activity of Hsp70 is shown in grey. Both DNAJA2 constructs activate the Hsp70 chaperone to the same extent. Data are means ± SEM (n = 3). **(I)** Refolding of heat-denatured firefly luciferase by a chaperone refolding system consisting of Hsp70, NEF, and WT DNAJA2 (red) or mutant DNAJA2$^{\Delta BH}$ (violet). DNAJA2 lacking the β-hairpin shows high protein refolding yields. Data are means ± SEM (n = 5).

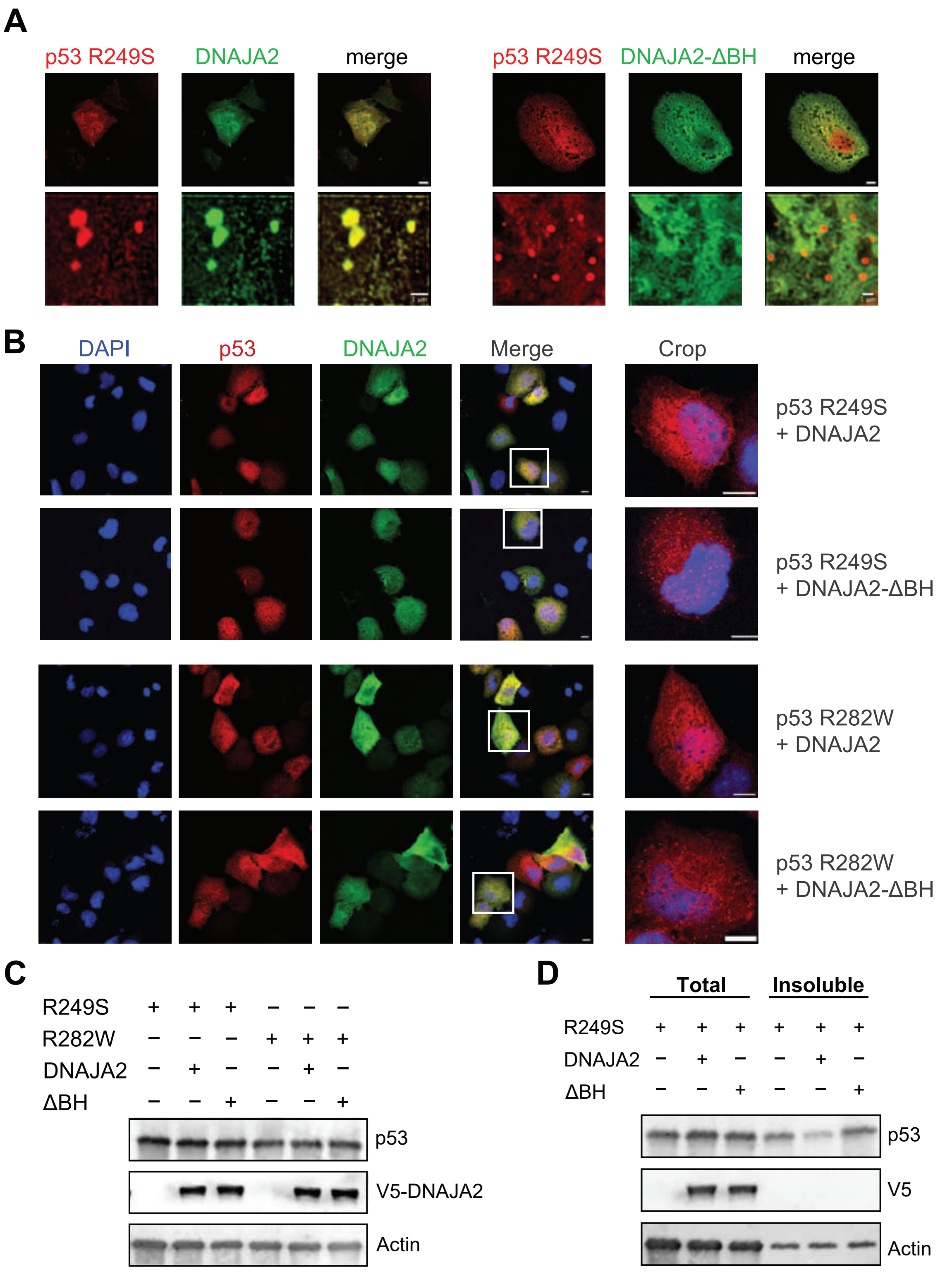

**Figure S5. DNAJA2 β-hairpin site is essential for p53 aggregation-prevention in cells. (A)** Colocalization of p53 R249S co-expressed with V5-DNAJA2 or V5-DNAJA2$^{\Delta BH}$ by stimulated emission depletion (STED) microscopy in SaOS2 cells. Scale bar: 10 μm and 1 μm as indicated. WT DNAJA2 co-localizes with residual p53 aggregates, while no co-localization is observed for DNAJA2$^{\Delta BH}$. The β-hairpin region thus drives DNAJA2 interaction with destabilized p53 in cells. **(B-C)** Immunostaining **(B)** and western blot of proteins levels **(C)** of mutant p53 (R249S, R282W) co-expressed with V5-DNAJA2 or V5-DNAJA2$^{\Delta BH}$, quantified in Figure 3F. The crop images (corresponding to white boxed areas in the merge image) are the overlay of the p53 signal with DAPI. Scale bar: 10 μm. The deletion of the β-hairpin region abolishes the prevention of aggregation activity of DNAJA2. **(D)** Supernatant/pellet fractionation of cell lysate of p53 R249S co-expressed with an empty vector, DNAJA2 and DNAJA2$^{\Delta BH}$. In contrast to WT DNAJA2, co-overexpression DNAJA2$^{\Delta BH}$ does not reduce the fraction of insoluble p53 mutant. Representative of n=3.

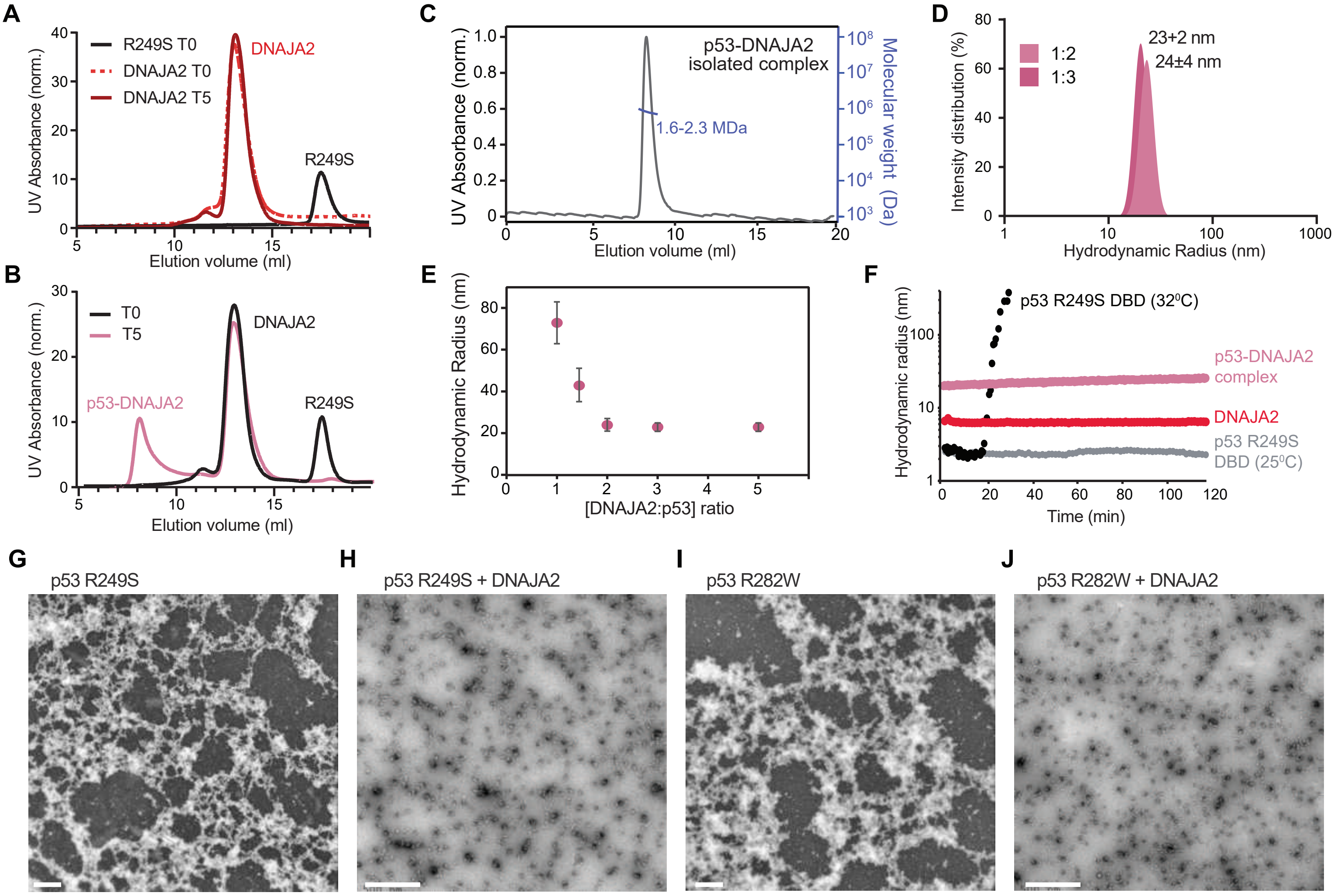

**Figure S6. DNAJA2 chaperones assemble into large oligomeric particles upon binding to misfolded p53. (A)** Size exclusion column (SEC) chromatograms of p53 (black) and DNAJA2 at time 0 (dark red) and following 5 hour incubation at 37°C (red dashed line). **(B)** SEC chromatograms following p53-DNAJA2 complex formation at time 0 (black), and following 5h incubation at 37°C (pink). **(C)** SEC-MALS analysis of isolated DNAJA2-p53 complexes separated on a Superdex 200 Increase 10/300 column. The DNAJA2-p53 complexes remain stable, with no visible dissociation of p53 monomers or DNAJA2 dimers. **(D)** Histograms of DLS-measured hydrodynamic radii of isolated R249S-DNAJA2 (1:2 and 1:3) complexes. **(E)** Hydrodynamic radius of DNAJA2-p53 complexes plotted as a function of DNAJA2 dimer concentration, as determined from DLS measurements. p53 concentration was kept constant at 3 μM. **(F)** DLS measurements of the hydrodynamic radius of isolated DNAJA2-p53 complex, p53 R249S DBD, or DNAJA2 measured for 2 hours at 32 °C. The hydrodynamic radius as a function of time for p53 R249S DBD under non-aggregation inducing conditions (25 °C) are shown as a control. The DNAJA2-p53 complex remains stable over time, while p53 R249S DBD without the DNAJA2 chaperone aggregates within 20 minutes. **(G-J)** Representative negative stain electron micrographs of p53 alone or p53-DNAJA2 mixtures following 3-hour incubation at 37°C. Both R249S **(G)** and R282W **(I)** p53 mutants form large amorphous aggregates in the absence of DNAJA2 and monodisperse ~40 nm particles when in complex with the chaperone **(H, J)**. Scale bar of 500 nm is shown on the bottom of each image.

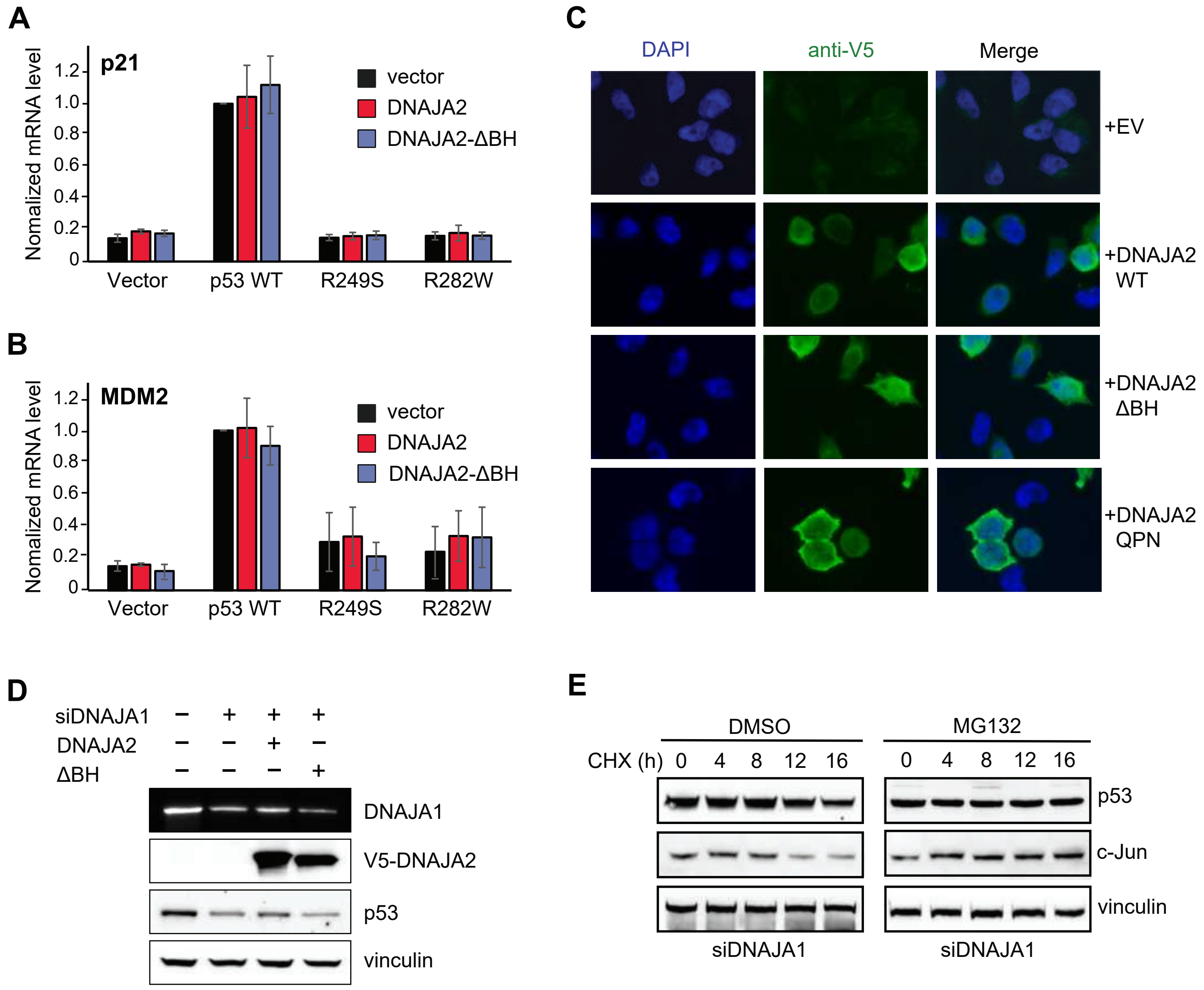

**Figure S7. Class A JDP chaperones protect the oncogenic p53 mutants from degradation. (A-B)** Quantification of mRNA levels of p53 target genes, p21 **(A)** and MDM2 **(B)**, in SaOS2 cells co-expressing p53 variants (WT, R249S, R282W) and V5-DNAJA2 or V5-DNAJA2$^{\Delta BH}$ by qPCR. Data represent mean values ± s.d (n=3). **(C)** Immunostaining of p53 and DNAJA2 in PaTu-8988 cells transfected with DNAJA1 siRNA and expressing DNAJA2$^{WT}$, DNAJA2$^{\Delta BH}$, or DNAJA2$^{QPN}$ variants. **(D)** Immunoblot monitoring p53, DNAJA1, DNAJA2, and vinculin levels in PaTu-8988 cells expressing empty vector, DNAJA2$^{WT}$, DNAJA2$^{\Delta BH}$ or DNAJA2$^{QPN}$ mutants with or without knockdown of DNAJA1. **(E)** MG-132 proteasome inhibitor blocks the turnover of endogenous p53 R282W protein, induced by the depletion of class A JDPs (siDNAJA1). Time points following treatment with cycloheximide (CHX) are shown on top. Vinculin and c-jun were monitored as controls.
